# Genome structure mapping with high-resolution 3D genomics and deep learning

**DOI:** 10.1101/2025.05.06.650874

**Authors:** Clarice K.Y. Hong, Fan Feng, Varshini Ramanathan, Jie Liu, Anders S. Hansen

**Affiliations:** Department of Biological Engineering, Massachusetts Institute of Technology; Cambridge, MA 02139, USA; The Broad Institute of MIT and Harvard; Cambridge, MA 02139, USA; Koch Institute for Integrative Cancer Research; Cambridge, MA, 02139, USA; The Novo Nordisk Foundation Center for Genomic Mechanisms of Disease, Broad Institute of MIT and Harvard, Cambridge, MA 02142, USA; Gilbert S. Omenn Department of Computational Medicine and Bioinformatics, University of Michigan, Ann Arbor, MI 48109, USA; Department of Computer Science and Engineering, University of Michigan, Ann Arbor, MI 48109, USA

**Author notes:** Co-corresponding authors: J.L. and A.S.H. These authors contributed equally to this work (Clarice K.Y. Hong, Fan Feng).

## Abstract

Gene expression is often regulated by distal enhancers through cell-type-specific 3D looping interactions, but comprehensive mapping of these interactions across cell types is experimentally intractable. To address this gap, we introduce an integrated approach where we generate ultra-deep Region Capture Micro-C (RCMC) and Micro-C data specifically designed for state-of-the-art deep learning architectures. We developed Cleopatra, an attention-based deep learning model that takes epigenomic inputs and is pre-trained on genome-wide Micro-C data followed by fine-tuning with high-resolution RCMC data. Cleopatra accurately predicts 3D maps at sub-kilobase bin sizes and unprecedented resolution, enabling us to generate ultra-high-resolution, genome-wide 3D contact maps across four human cell types. These maps revealed cell-type-specific microcompartments and over 900,000 loops across the cell types, about half of which are cell-type-specific. Using Cleopatra maps, we observe that promoters form about a dozen loops on average, and that expression increases monotonically with the number of loops, indicating that looping is associated with higher gene expression. We further show the enhancer-promoter loops are often anchored by CTCF, and nominate new transcription factors that may regulate cell-type-specific enhancer-promoter interactions. Overall, we establish a framework for ultra-high-resolution 3D genome mapping, providing a broadly applicable resource for gaining new insights into cell-type-specific gene regulation.

## Introduction

Precise regulation of gene expression is essential for cellular function, and disruption of gene expression is a major cause of disease. Genes are often regulated by distal enhancers that loop to their cognate promoters in a cell-type specific manner to drive cell-type-specific gene expression^1^, sometimes up to more than a megabase away. Since the human genome is thought to contain ∼ 20,000 genes and hundreds of thousands of enhancers^2^, and enhancers sometimes skip nearby genes to regulate more distal genes^3–6^, understanding cell-type-specific enhancer-promoter (E-P) interactions and gene regulation is quite challenging^2^. Thus, comprehensive 3D genome maps that exhaustively capture all loops in multiple cell types are required to understand the principles of distal gene regulation in human health and disease.

The 3D genomics method Hi-C has transformed our understanding of 3D genome folding^7–12^. While Hi-C resolves A/B-compartments, topologically associating domains (TADs), and structural CTCF/cohesin loops^10^, Hi-C fails to detect most loops between *cis*-regulatory elements (CREs)^13,14^. Micro-C overcomes this limitation by effectively detecting CRE loops in addition to the larger scale structural features seen in Hi-C^13–15^, but mapping of all 3D interactions across the human genome at nucleosome-scale bin sizes is intractable due to the extreme sequencing required to cover all pairwise interactions. To address this issue, we previously developed Region Capture Micro-C (RCMC)^16^ to capture selected megabase-sized regions from Micro-C libraries, enabling us to generate 3D genome maps at unprecedented resolution with dramatically less sequencing. While RCMC maps revealed fine-scale CRE loops and microcompartment structures at small bin sizes that were not previously visible in Micro-C, it typically only captures *<* 0.5% of the genome. It is therefore not experimentally feasible to map fine-scale 3D genome structure and all chromatin loops across the entire genome.

Instead, several deep learning models have been developed to predict 3D genome structure. These models can robustly predict Hi-C maps and large-scale structures at kilobase to megabase scale such as A/B-compartments, TADs, and CTCF loops^17–29^. However, no deep learning model can currently predict 3D genome maps with sufficient detail to identify fine-scale genome structure and comprehensively map CRE loops genome-wide.

Here, we overcome these challenges by integrating experimental Micro-C and RCMC with a new attention-based deep learning model, Cleopatra. We generate genome-wide Micro-C maps and ultra-deep RCMC maps for 14 diverse regions in four human cell lines to capture cell-type-specific differences in 3D genome structure. This enables a pre-training/fine-tuning approach, where we pretrain Cleopatra on genome-wide Micro-C with 19 different genomic features, fine-tune on RCMC, then predict ultra-high-resolution genome-wide 3D maps that capture finescale genome structures. Our integrated experimental-deep learning approach allowed us to generate the most comprehensive map of 3D genome structure and loops across the human genome and begin to unravel the general principles of distal gene regulation.

## Results

### Ultra-high-resolution 3D genome maps in four human cell types

To train Cleopatra, we first generated the deepest 3D contact maps in human cells to date. We performed RCMC in 14 regions across four cell types, GM12878 (lymphoblastoid), HCT116 (colorectal carcinoma), K562 (chronic myelogenous leukemia), and H1 (human embryonic stem cells, hESCs) (Figure 1A). The cell types were selected to be diverse and have been extensively profiled by the ENCODE and Roadmap Epigenomics consortia^30–33^. We selected 2-3Mb regions covering diverse epigenomic signals, gene densities, and expression levels (Figure S1). We sequenced each region to an average of 95 billion genomewide equivalent reads in all cell types (Methods, Table S1), with the most deeply sequenced regions in GM12878 exceeding 500 billion genome-wide equivalent reads. We also generated Micro-C maps for GM12878, HCT116, and K562 with ∼5-7 billion unique pairwise contacts.

**Figure 1:**
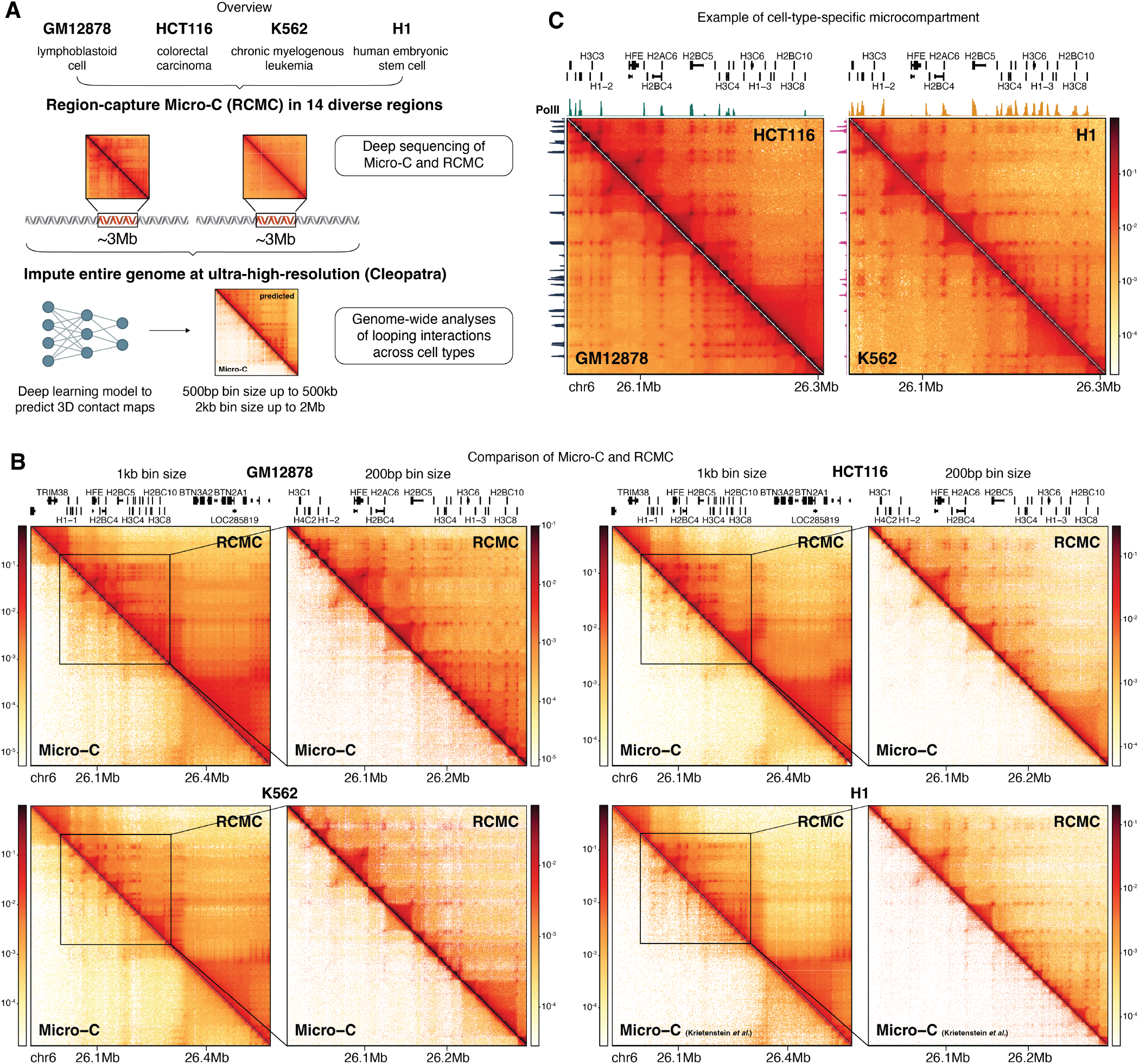
RCMC enables ultra-high-resolution 3D genome mapping and boundary identification. (A) Overview of study workflow. We first performed genome-wide Micro-C and RCMC in 14 regions across four cell types, which was then used to train Cleopatra to predict ultra-high-resolution 3D contact maps for the entire genome. (B) Representative contact maps (region1) comparing RCMC to best available Micro-C data in four cell types. (C) Example of cell-type-specific microcompartment, where interactions in each cell type are correlated with RNA Pol II binding and transcription. POLR2A ChIP-seq tracks for the corresponding cell type are plotted on the left for the map on the bottom left and on top for the map on the top right. Contact maps are plotted at 1kb bin size.

To validate our maps, we first show that our Micro-C and RCMC maps correspond well with existing Hi-C data^10^ at coarse bin sizes, including A/B-compartments, TADs and structural CTCF/cohesin loops (Figures S2A-S2B). Our RCMC data also shows the same expected distance decay as Micro-C (Figure S2C). However, at small bin sizes, RCMC far outperforms current Hi-C and Micro-C datasets both visually and quantitatively (Figures 1B, S2D-S2G). Using the definition of resolution in Rao, Huntley *et al*.^8^, our RCMC maps achieved 50bp-100bp resolution for most regions (Table S1). However, resolution is also commonly used to refer to “ bin size” . To avoid confusion, we use resolution to refer to how well we can resolve fine-scale structures (higher resolution means more resolved structures) and bin size when referring to contact map data. In total, we generated ultra-high-resolution 3D maps over ∼ 44Mb in each of the four human cell types.

Using RCMC, we previously discovered microcompartments in mouse embryonic stem cells (mESCs), a pattern of highly nested and focal genome interactions that can only be robustly detected in RCMC but not Micro-C or Hi-C data^16^. Given that we only profiled one mouse cell type, it was unclear whether microcompartments are cell-typespecific genome structures. Here we show that microcompartments also form in human cells and can be cell-type specific. For example, the histone replication-dependent genes form microcompartments in all cell types, but the exact loop anchors within the compartment depends on which genes are transcribed (Figure 1C). In GM12878, the essential B-cell development gene *EBF1* ^34^ forms a GM12878-specific microcompartment (Figure S3A). Microcompartment anchors are not simply an artifact of open chromatin regions, as not all interactions are anchored by DNase-accessible peaks (Figure S3B; Supplementary Note 1)^16^. Thus, while many larger-scale A/B-compartments and TADs appear to be largely consistent across cell types^8^, smaller-scale microcompartments are cell-type-specific 3D genome structures that may be associated with cell-type-specific transcription.

### RCMC enables accurate identification of cell-type-specific boundaries and loops

We next asked if the fine-scale boundaries associated with microcompartment anchors are similarly cell-type specific. Micro-C previously revealed boundaries at 200bp bin sizes that cannot be detected in Hi-C^13^. Applying a similiar analysis to RCMC, we find that although RCMC maps are much deeper than Micro-C, we identified fewer boundaries in RCMC compared to Micro-C (Figure S4A). Visual inspection suggested that boundary calling in Micro-C is prone to false positives due to lower depth and sampling noise (Figure S4B). We confirmed this by showing that there are more boundary calls in downsampled RCMC maps (Figure S4C), and that known boundary-associated features such as CTCF, PolII and H3K27ac are enriched at RCMC-specific boundaries and depleted at Micro-C-specific boundaries (Figure S4D). With more accurate RCMC boundaries, we find that fine-scale boundaries tend to be cell-type specific, with ∼ 50% of boundaries found in only one cell type (Figure 2A). Boundaries that are shared among cell types are enriched for CTCF motifs and tend to be stronger, while cell-type-specific boundaries are enriched in motifs for cell-type-specific transcription factors (TFs) and are weaker (Figures 2B & S4E). For example, GM12878-specific boundaries are enriched for IRF motifs, which are a class of important B-cell development TFs^35^,^36^, while K562 boundaries are enriched for GATA motifs, which are crucial for blood development^37^. These results emphasize the need for high resolution RCMC maps to accurately identify fine-scale boundaries of small domains and microcompartments, enabling us to nominate new TFs as 3D genome regulators that may contribute to the formation of cell-type-specific 3D structures.

**Figure 2:**
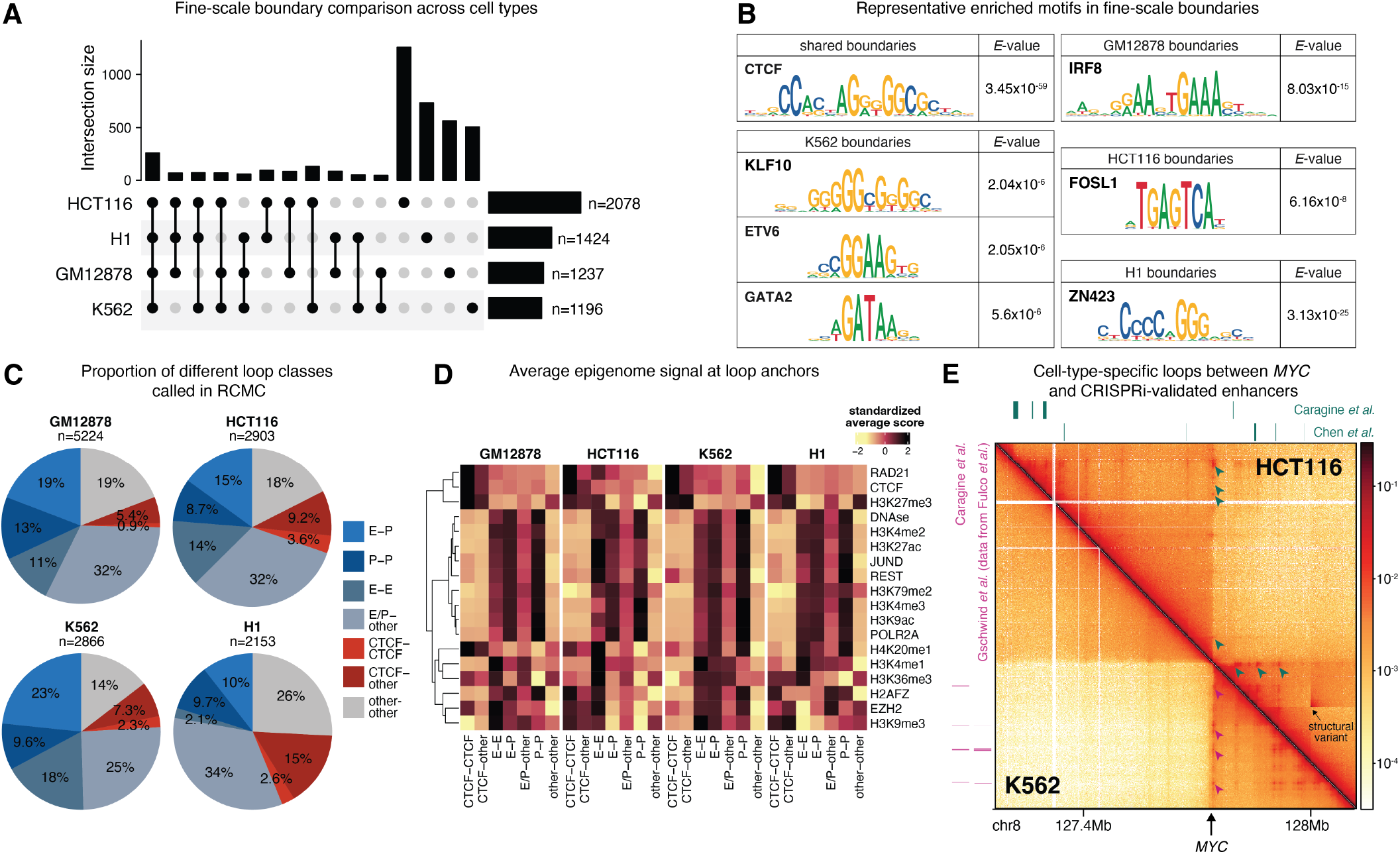
Ultra-deep RCMC reveals cell-type-specific microcompartments and CRE interactions. (A) Comparison of fine-scale boundary locations across all cell types. n indicates the number of boundaries called in each cell type. (B) Representative motifs enriched in boundaries that are shared between cell types or specific to each cell type. *E* -values were computed by AME^82^. Additional motifs can be found in Table S2. (C) Loops called in RCMC are enriched for CRE interactions such as E-P, P-P and E-E loops. n indicates the number of loops called in each cell type and includes both manually annotated and CHIRON-called loops. (D) Mean signals of various ChIP signals for each loop class. Coverage of each signal was calculated for each 1kb loop anchor, then averaged across all loop anchors in a given loop class. The signals were then standardized across the row to ensure that all signals are on the same scale. (E) RCMC detects cell-type-specific and CRISPRi validated looping interactions with *MYC* (indicated at bottom). Bars represent validated enhancers identified by the respective citations (Caragine *et al*.^43^, *Gschwind et al*.^54^, *Chen et al*.^46^*). Caragine et al*. enhancers for HCT116 are from HT29, which is a different colorectal cancer cell line. Circled loops correspond to *MYC* looping interactions that match the respective validated enhancers. The contact map is plotted at 1kb bin size.

RCMC also enables the identification of more fine-scale loops than Micro-C^16^. Using a combination of manual annotations and a new deep learning-based loop caller, CHIRON (Figures S5-S6, Supplementary Note 2), we identified 5,224 loops in GM12878, 2,676 loops in HCT116, 2,866 loops in K562 and 2,154 loops in H1 cells in RCMC data at 1kb bin size. The larger number of loops in GM12878 is mainly due to deeper sequencing, as downsampling GM12878 reduces the number of loops called (Figure S5F). Similar to prior studies, we observe that the loops are enriched for *cis*-regulatory interactions, including E-P, P-P and E-E interactions^13^,^16^ (Figure 2C). However, about half of the loops have one unannotated anchor (“ other”) and about a quarter are not annotated on both anchors (“ other-other” ; Figure 2C). To gain insight into the identity of these unannotated anchors, we quantified common epigenomic features at loop anchors (Figure 2D). As expected, CRE loop anchors (E-P, P-P, and E-E) are enriched for active modifications such as H3K4me1 and H3K27ac, whereas CTCF-CTCF interactions are enriched for CTCF and RAD21. Other-other loops are enriched for H3K27me3 and/or H3K9me3, consistent with a putative role for generally repressive marks in loop formation^38^,^39^. We also did not observe strong overlap with previously reported silencers in K562 cells^40^,^41^. Approximately half of all loops in each cell type are cell-type specific, with cell-type-specific loops spanning all loop annotations and enriched in “ other-other” loops (Figure S7). Overall, RCMC enables comprehensive identification of CRE loops, though many cell-type-specific loops fall into the “ other” category that is not well characterized by known epigenomic factors. This suggests that there may be other classes of looping interactions beyond CTCF and CRE loops that remain uncharacterized.

In theory, other capture-based methods such as promoter capture Hi-C^42^ or ChIA-PET/HiChIP^4^,^43^ can be used to identify genome-wide loops without extensive sequencing. However, while RCMC can be rigorously normalized using matrix balancing^16^,^44^, robust normalization of capture-based methods is more challenging (Supplementary Note 3). We also found that promoter-capture Hi-C, RNA Pol II ChIA-PET and H3K27ac HiChIP were prone to false positive or false negative loop detection depending on the study, and that the strength of shared loops show poor correlation between RCMC and promoter-capture Hi-C (Figure S8, Supplementary Note 3). To investigate if the RCMC-detected loops are functional, we compared RCMC data against CRISPRi datasets at the *MYC* region in K562 and HCT116^3,43,45,46^. RCMC detects cell-type-specific *MYC* interactions corresponding only to the enhancers validated in each cell type (Figure 2E). Together, we show that RCMC detects cell-type-specific and functional looping interactions.

### Cleopatra predicts 3D genome structure at ultra high resolution

While RCMC enabled us to generate ultra-high resolution 3D genome maps, it only provided information for ∼ 44Mb of the whole genome. To extend the resolution of RCMC to the entire genome, we developed Cleopatra, a deep learning model that accurately predicts whole-genome contact maps at the resolution of RCMC (Figure 3A). We had previously developed CAESAR to predict Micro-C contact maps at nucleosome-sized bins^24^, but its architecture and learning strategy are insufficient to predict RCMC maps (Supplementary Note 4). Cleopatra uses 19 cell type-specific genomic features as input (DNase-seq, H3K4me1, H3K4me2, H3K4me3, H3K9ac, H3K9me3, H3K27ac, H3K27me3, H3K36me3, H3K79me2, H4K20me1, CTCF, EZH2, POLR2A, JUND, REST, RAD21, H2AFZ, and phastCons score), and predicts contact maps at two bin sizes (500 bp and 2kb). To learn local genomic patterns and capture chromatin states, Cleopatra contains three convolutional blocks. To further capture long-range dependencies corresponding to looping interactions between distal elements, we included state-of-the-art self-attention blocks, which are at the core of the Transformer structure^47^ and the basic unit of modern language models^48,49^.

**Figure 3:**
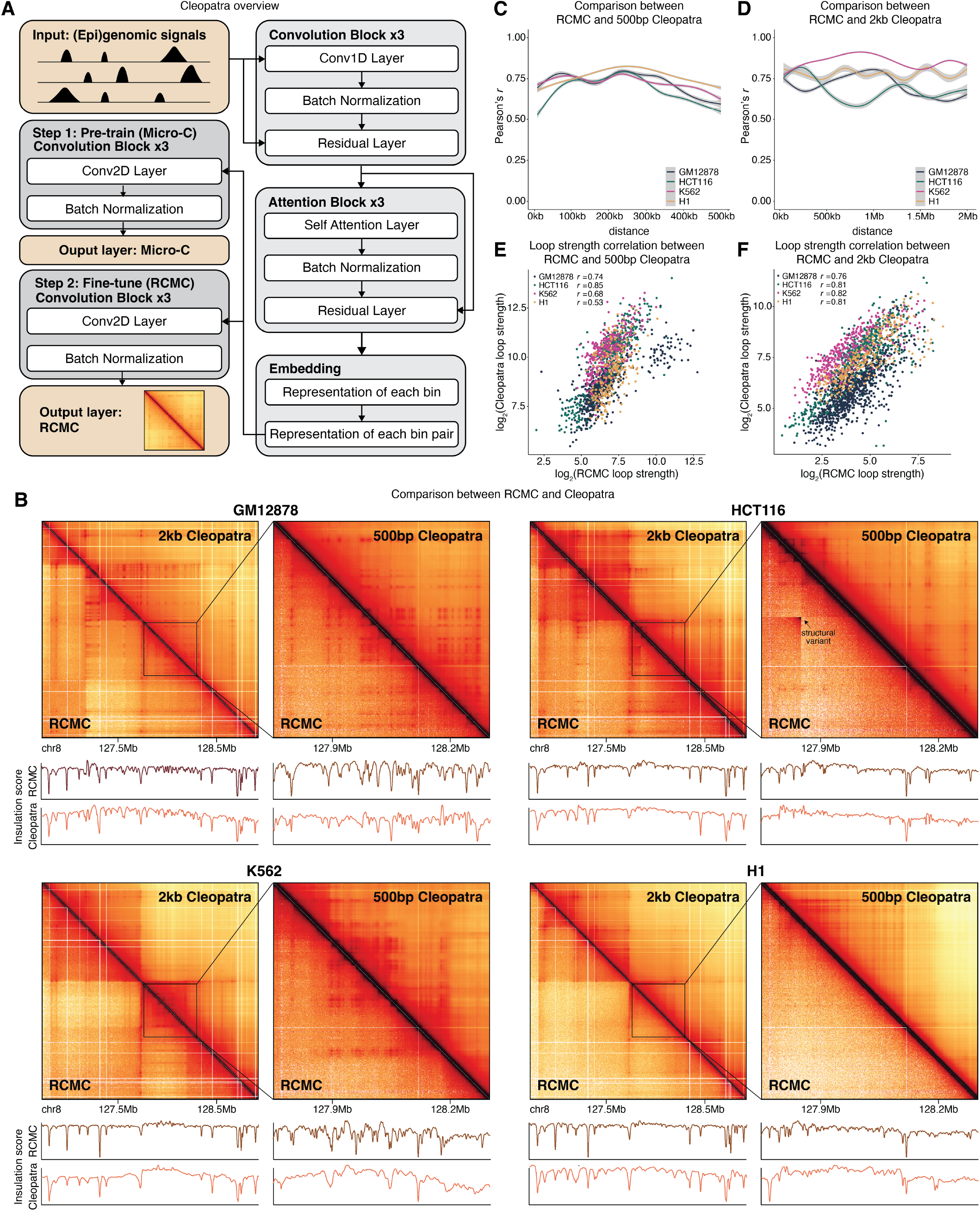
Cleopatra predicts ultra-high-resolution contact maps for four human cell types. (A) Overview of Cleopatra method. Epigenomic signals for each cell type are first pre-trained with Micro-C maps before fine-tuning with RCMC. (B) Cleopatra predictions in a representative holdout region (region6) across four cell types at 2kb (left) and 500bp (right) bin sizes. Insulation scores for the corresponding regions from RCMC and Cleopatra are plotted below the maps. (C)-(D) Distance stratified pearson’s correlations between RCMC and 500bp (C) or 2kb (D) Cleopatra model in three holdout regions. Shaded area represents 0.95 confidence interval from loess smoothing. (E)-(F) Loop strength correlation between RCMC and 500bp (E) or 2kb (F) Cleopatra models using loops called in RCMC. *r* represents Pearson’s correlations for each cell type.

We trained individual Cleopatra models at 500bp and 2kb bin size for each cell type. All contact maps were converted to log(observed/expected), and the genome was segmented into windows of 1,250 bins (625kb for 500bp bin size and 2.5Mb for 2kb bin size), with each window serving as an individual training example. During prediction, Cleopatra outputs contact maps using a sliding window of 1,250 bins with a 250-bin step length, which effectively covers interactions within 1,000 bins (500kb for 500bp bin size and 2Mb for 2kb bin size). Finally, the windows are merged to generate continuous predictions across the genome, and the observed/expected pixel values were scaled to “ observed” values to facilitate comparisons with experimental data (Methods). Cleopatra takes full advantage of our experimental data with a two-stage training strategy. The model is first pre-trained on wholegenome Micro-C data, then fine-tuned on much deeper RCMC data, allowing Cleopatra to effectively transfer knowledge from more abundant but lower-resolution data to make accurate, high-resolution genome-wide contact map predictions. During pre-training, we aimed to minimize mean squared errors (MSEs) between experimental and predicted maps. In the fine-tuning stage, to better capture the fine-scale domains and microcompartments in RCMC, we integrated the MSEs of insulation scores into the loss function and added additional weights on loop regions. Ultimately, for each cell type, we generate 500bp bin size predictions for interactions up to 500kb and 2kb bin size predictions for interactions up to 2Mb.

To evaluate the models, ten regions were used to finetune Cleopatra and three regions (regions 4, 6 & 11) were held out to compare with RCMC (Methods). Cleopatra accurately predicts highly detailed, dense 3D contact maps that strongly resemble RCMC contact maps (Figures 3B & S9A), achieving high Pearson’s correlation coefficients in holdout regions across different prediction distances (Figures 3C-3D). Both 500bp and 2kb predicted maps also agree well with each other (Figures S9B-S9C). Loop strengths in RCMC and Cleopatra maps correlate well in all cell types, indicating that Cleopatra accurately predicts quantitative loop strengths in holdout regions (Figures 3E-3F). Compared to other deep learning 3D genome models^17–29^, Cleopatra is unique in using a pre-training/fine-tuning approach. To evaluate the importance of fine-tuning on RCMC data, we compared the output of the pre-training with the fine-tuned model. The pre-trained model cannot predict RCMC-specific structures at 500bp bin sizes (Figures S10A-S10B), suggesting that the model learns RCMC-specific structures during the fine-tuning step despite the limited number of genomic regions. However, pre-trained 2kb Cleopatra appears sufficient to predict coarser RCMC-like maps (Figures S10C-S10D), likely because kilobase-scale structures are also weakly visible in Micro-C maps. These results emphasize the need for RCMC to develop models that can predict highly detailed 3D contact maps at sub-kilobase bin sizes. To facilitate ease of use by the community, we provide Cleopatra predictions in Cooler format^50^, allowing users to easily view Cleopatra maps of any region of interest.

Notably, Cleopatra predicts contact maps at smaller bin sizes than most existing deep learning models for predicting 3D genome structure^51^, including Akita^17^, HiCPlus^18^, Epiphany^20^, and C. Origami^21^, which were designed for coarser bin sizes (typically 2–10kb). For example, the most recent Akita v2 model predicts at 2048bp bin sizes up to 1Mb distance^29^, while ChromaFold and C.Origami predict up to 2Mb distances at 10kb and 8192bp bin sizes respectively^21^,^28^. Comparing Cleopatra to Akita v2 and C.Origami, we show that all models can predict large-scale structures such as TADs, but only Cleopatra predicts the fine-scale interactions unique to RCMC (Figure S11). The contact maps predicted by Cleopatra are therefore uniquely suited for fine-scale analyses of *cis*-regulatory interactions across the genome.

The full version of Cleopatra uses 19 different input features to maximize performance for the common cell types used here. To enable more widespread adoption of the Cleopatra framework, we asked if a smaller set of features is sufficient to predict fine-scale features. Based on the correlation of features, availability of features in the ENCODE dataset and features used in prior deep learning models^20,21^, we selected features to run models with 7, 5 and 3 inputs respectively in GM12878, which has the best RCMC data and predictions (Figure S12A). We show that model performance decreases slightly as the number of features are reduced (Figures S12B-S12C), and fine-scale loops and microcompartments become increasingly blurry with fewer inputs in some regions (Figures S12D-S12E). Similarly, loop strength correlations slightly decrease with fewer inputs, though the effect is more pronounced in the 2Mb model (Figures S12F-S12G) However, we note that both 500kb and 2Mb 7-input models perform almost as well as the full model. We trained 7-input models for all cell types, and indeed observed similar performance for all cell types (Figures S12H-S12I). Thus, we show that a 7-input core set of features (DNase-seq, H3K4me1, H3K4me3, H3K9me3, H3K27ac, H3K27me3, and CTCF) can be sufficient for accurate predictions, though we note the number of inputs may be further reduced depending on the application.

To delineate the contribution of each feature, we asked how loop predictions in Cleopatra maps change with fewer inputs. Because the role of CTCF in 3D looping is most well documented, we trained 7-input models without CTCF in GM12878. As expected, dropping CTCF leads to a decrease in model performance, going below even the 3-input model for 500bp Cleopatra (Figures S13A-S13B), reducing Cleopatra’s ability to predict CTCF-anchored loops (Figure S13C). In contrast, loss of H3K4me1 from the 7-input to 5-input model does not reduce the average loop strength of H3K4me1-anchored loops, while the loss of H3K4me3 from the 5-input to 3-input model only mildly reduces average loop strength of H3K4me3-anchored loops (Figures S13D-S13E), even though the overall performance of the 5-input and 3-input loops is reduced. We conclude that while CTCF is the primary feature driving CTCF-anchored loops in Cleopatra as expected, most other loops are predicted from a complex interplay of multiple features.

### Cleopatra reveals cell-type-specific micro-compartments and loops across the genome

We next used our ultra-high-resolution Cleopatra maps to analyze microcompartments and loops across the entire genome. We identified new microcompartments across the genome that are not visible in Micro-C data (Figures 4A, S14A), including several regions where cell-type-specific microcompartments are predicted (Figure 4B), highlighting the utility of Cleopatra in identifying cell-type-specific 3D genome structures.

**Figure 4:**
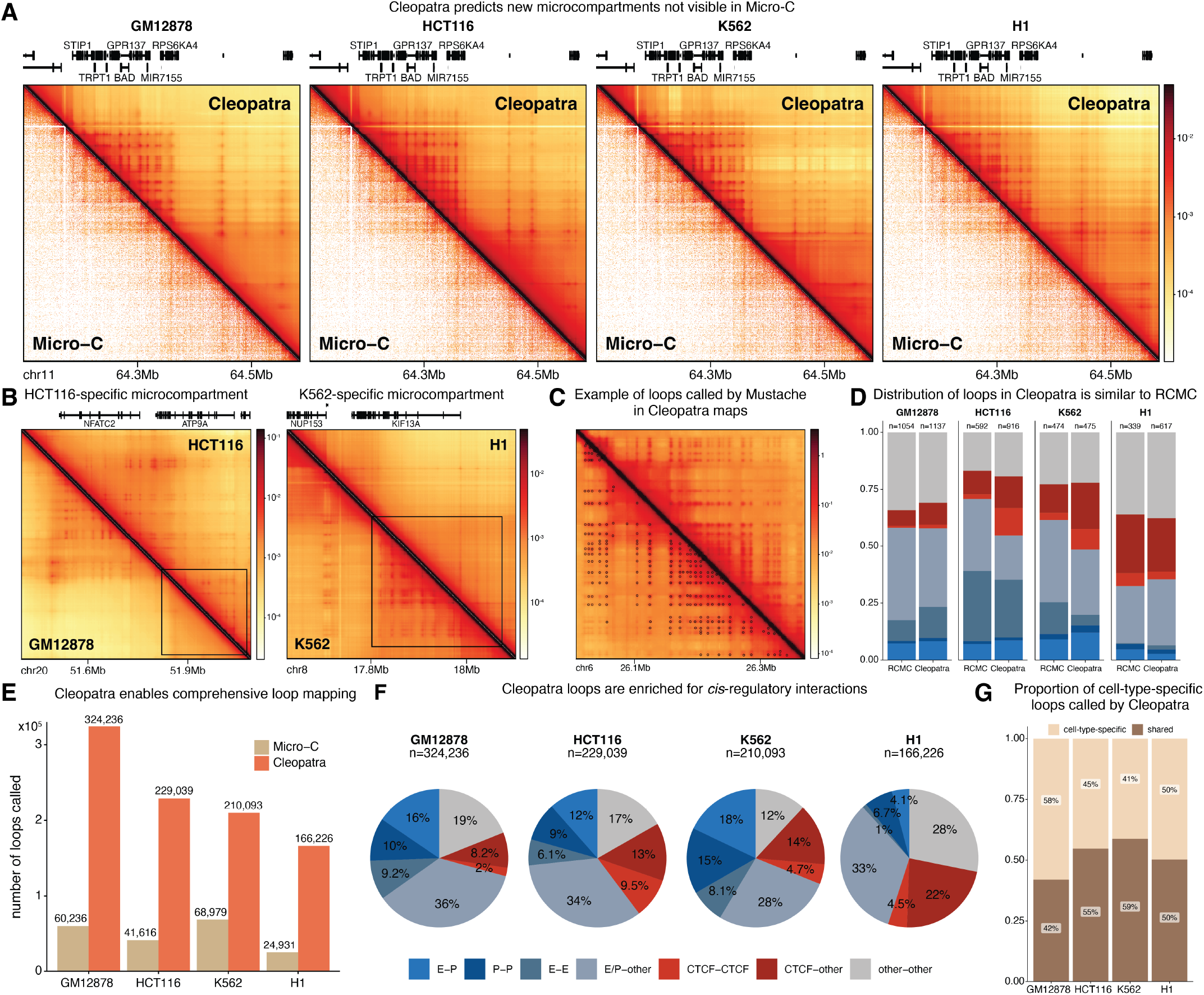
Cleopatra reveals new cell-type-specific microcompartments and loops. (A) Representative new microcompartment predicted by Cleopatra (500bp model) that is not visible in Micro-C. (B) Example cell-type-specific microcompartment (box) in HCT116 (left) and K562 (right) predicted by Cleopatra (500bp model). (C) Example of loop calls by Mustache in Cleopatra contact map (500bp model). (D) Distribution of loop classes in Cleopatra holdout regions compared to RCMC. Colors represent loop classes as in (F). (E) Number of loops called by Mustache in Cleopatra-predicted maps and Micro-C. (F) Loop class distribution for all Cleopatra predicted loops. n indicates number of loops for each cell type. (G) Proportion of predicted loops that are shared with at least one other cell type (shared) or only present in the respective cell type (cell-type-specific).

To call loops in Cleopatra maps, we used Mustache^52^ (Figure 4C) and merged the loops from the 500bp and 2kb predictions, using the 500bp loop calls for distances *<*495kb and 2kb loop calls for distances between 495kb and 1.98Mb. In the three holdout regions, we found similar numbers of loops as RCMC, with a similar distribution of loop classes (Figure 4D), demonstrating that Cleopatra detects the full spectrum of loops experimentally observed in RCMC data. Across the genome, we identified 324,236 loops in GM12878, 229,039 loops in HCT116, 210,093 loops in K562, and 166,226 loops in H1 predicted maps. This is about an order of magnitude more loops than previously identified in the best available Micro-C maps (22,508 loops in H1 and 36,989 loops in human foreskin fibroblast cells)^14^ and the Micro-C maps generated in this study (Figure 4E). Cleopatra predicts the majority of CRE loops called in Micro-C, but performs less well on other-other loops, suggesting that other-other loops are more difficult to accurately predict (Figure S14B). This is likely because “ other” anchors are not well-marked by most known epigenomic signals. Nevertheless, this observation indicates that Cleopatra is not simply overfitting on Micro-C maps, but is learning how the input epigenomic features relate to *cis*-regulatory looping. Indeed, classification of loop anchors reveals that most loops are *cis*-regulatory loops (Figure 4F) with a median loop size of ∼ 200-300kb (Figure S14C), showing that Cleopatra predicts novel interactions between regulatory elements across the entire genome. The predicted loop anchors share similar characteristics to RCMC loop anchors, with other-other loops also enriched for H3K27me3 signals (Figure S14D). Next, we investigated how TADs may influence looping interactions. In GM12878, which has the largest number of loops predicted by Cleopatra, we find that 233,385/324,236 (72%) of all loops and 38,665/50,313 (77%) of E-P loops cross TAD boundaries, suggesting that they might be insufficient to fully restrict most looping interactions^53^. Finally, we asked if Cleopatra loops are cell-type-specific. Similar to RCMC analyses, about 40-60% of loops are unique to each cell type, and cell-type-specific loops are enriched for E/P-other or other-other interactions (Figures 4G & S14E). Taken together, we show that Cleopatra predictions are an important resource for identifying new CRE interactions.

To validate that Cleopatra predicts functional looping interactions, we tested the correspondence between Cleopatra loops and a list of harmonized, high-confidence regulated E-P pairs curated by the ENCODE consortium in K562 cells^54^. We find that high-confidence “ regulated” E-P pairs have stronger loops than the non-regulated E-P pairs (Figure S14F). We also identified a set of high-confidence expression quantitative trait loci (eQTLs) in GM12878^55^ and show that eQTL-promoter interactions are strongest in GM12878 (Figure S14G). Thus, using Cleopatra maps, we have generated the most comprehensive set of cell-type-specific looping interactions in the human genome across four cell types.

### Functional analyses of genome-wide *cis*-regulatory interactions

The comprehensive Cleopatra loop calls enabled us to investigate the relationship between looping and gene regulation at fine scales. We find that while 86-92% of expressed genes make loops, only 21-39% of inactive genes make loops (Figure S15A). Overall, promoters form a median of 3-9 loops depending on cell type, with expressed promoters forming 5-18 loops (Figure 5A). We asked whether gene expression levels are generally controlled by single dominant enhancers or by the weak contribution of many enhancers. We observed that gene expression increases largely monotonically with the number of loops to the promoter, though adding more loops eventually results in diminishing returns (Figure 5B). A single E-P or P-P loop is associated with high gene expression, with additional loops increasing expression to a smaller extent, while P-other loops are correlated with a more gradual increase in loop expression (Figures 5C & S15B). From the enhancer point of view, the majority of annotated enhancers form E-P loops (GM12878: 83%, HCT116: 77%, K562: 75%, H1: 68%) with a median of 1-3 promoters per enhancer (Figure 5D). An open question in E-P looping is whether enhancers generally loop to the closest gene^56,57^ or whether they skip the closest gene to loop to more distal genes^3–5^. Using Cleopatra predicted loops, we find that about one-third of all enhancers (GM12989: 13,617, HCT116: 7,404, K562: 12,440, H1: 3,560) skip the nearest gene to loop to genes that are further away (Figure 5E). While skipped genes are enriched for inactive genes, the majority of skipped genes are still expressed, albeit at lower levels than the corresponding looped promoter (Figures S15C-S15D). Given the prevalence of enhancer skipping, it is important to exhaustively map all CRE-promoter interactions to understand the fundamental mechanisms of gene regulation and for many biological applications including functional interpretation of non-coding disease variants^6^.

**Figure 5:**
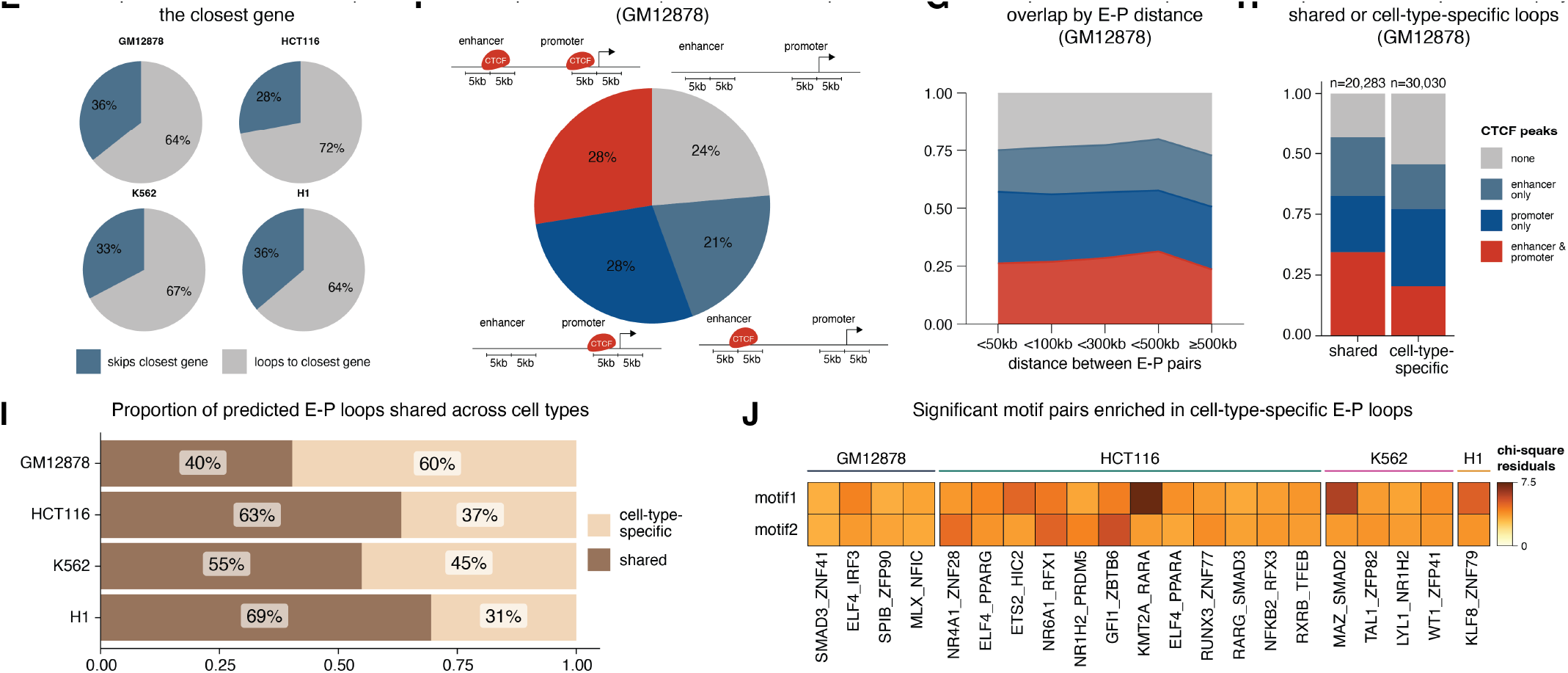
Functional analyses of looping interactions across the genome. (A) Number of looping interactions per promoter for all promoters or expressed promoters only. Each point represents a single promoter. (B) Median expression of promoters with increasing numbers of loops. Only promoters with *≤* 20 loops are shown here. (C) Median expression of promoters with increasing numbers of E-P/P-P/P-other loops in GM12878. Only promoters with *≤* 7 loops are shown here. In and (C), expression is calculated as the median FPKM of all promoters, including non-expressed promoters. (D) Number of E-P loops formed per enhancer. Each point represents a single enhancer. (E) Proportion of active enhancers that skip both their neighboring genes on both sides. (F) Proportion of E-P loops with CTCF binding at both the enhancer and promoter or enhancer/promoter only in GM12878. CTCF binding is defined using CTCF ChIP-seq peak calls. Colors correspond to legend in (H). (G) Proportion of E-P loops with CTCF binding on either anchor stratified by distance between the enhancer and promoter in GM12878. Colors correspond to legend in (H). (H) Proportion of E-P loops with CTCF binding on either anchor grouped by whether the E-P interaction is cell-type specific or shared with at least one other cell type in GM12878. (I)Proportion of cell-type-specific or shared E-P loops in each cell type. (J) Enrichment of significant motif pairs in cell-typespecific E-P loops. A chi-squared test was used to calculate the enrichment of each pair over loops that only contain one of the two motifs, resulting in two values for each motif pair. Motifs within the pair are arranged alphabetically, and the top row corresponds to enrichment of the pair over the first motif, while the bottom row corresponds to enrichment of the pair over the second motif. Only motif pairs with p-values of *<*0.05 over both motifs after Benjamini & Hochberg correction were considered significant and plotted here.

CTCF binding at CREs has been proposed to facilitate E-P interactions by increasing the strength of interactions between enhancers and promoters^1,5,58–62^. Using a 5kb window size around the CRE midpoint, we find that the four possible configurations (CTCF binding at promoter, enhancer, both, or neither) are about equally common (Figures 5F & S16A), with the contribution of CTCF further increasing if a more lenient window size is used (Figure S16B). Thus, Cleopatra predicts that the majority of human E-P loops appear to be at least partially facilitated by CTCF. Although CTCF is thought to be particularly important for long-range E-P loops^1,60,62,63^, the proportion of E-P loops facilitated by CTCF does not strongly increase with distance (Figures 5G & S16C). We also find that loops that are shared across cell types are more likely to have CTCF binding at either end, consistent with the observations made by Chen *et al*.^5^ (Figures 5H & S16D). While CTCF depletion experiments will be required to understand the effects on transcription, these results suggest that about 3/4 of E-P loops might be at least partially facilitated by CTCF in a largely distance-independent manner.

Housekeeping genes are thought to be primarily regulated by proximal enhancers, while developmental genes are regulated by more distal enhancers^64,65^. Yet, to our knowledge, 3D interactions of both types of promoters have not been systematically characterized. Using Cleopatra predictions, we find that E-P distances for enhancers and promoters are similar, suggesting that developmental enhancers may not be more distally located than housekeeping enhancers (Figure S17A). Housekeeping promoters also form more loops than developmental promoters on average, even when only accounting for E-P loops (Figures S17B-17C). We next asked if developmental promoter interactions are more cell-type specific. We find that approximately 50% of loops are cell-type-specific for both classes of promoters, suggesting that even housekeeping promoters may be regulated by different enhancers in different cell types (Figure S17D). For example, the housekeeping gene ZNF143’s^66–68^ promoter forms a unique E-P loop in GM12878 that is not present in other cell types (Figure S17E). Our results suggest that the enhancers that regulate housekeeping and developmental genes may be more similar than previously suggested.

Cell-type-specific loops are crucial for maintaining cellular identity and thought to be partially mediated by TFs^69–74^. We therefore sought to identify features of celltype-specific E-P loops. Of all E-P loops in each cell type, 30-60% are cell-type-specific (Figure 5I). However, the TFs that regulate cell-type-specific interactions are still unknown. Taking advantage of the large number of loops predicted by Cleopatra, we identified motifs at E-P anchors and searched for enrichment of motif pairs at cell-type-specific vs shared E-P loops (Figure 5J). In all cell types, we identified only a small number of statistically significant motif pairs. SMAD and ELF motifs were identified in multiple cell types but associate with a unique partner in each cell type, suggesting that the same TFs may mediate cell-type-specific loops by interacting with different partners in different cell types. In K562, we identified a SMAD2-MAZ pair, which has previously been found to mediate loops with CTCF^75^ and putatively with MAX/MYC/CHD2^76^. TAL1 has also been previously found to regulate chromatin loops at the globin locus along with LDB1, LMO2 and GATA1^77^, and here we further find that it may partner with ZFP82. In GM12878, we identified the ELF4-IRF3 pair, which are both TFs that regulate interferon expression and have been found to cooperatively interact at interferon gene promoters^78^. While perturbation experiments will be required for validation, these results illustrate how the large number of loops identified by Cleopatra allows us to nominate new interacting pairs that may regulate cell-type-specific E-P looping across the genome.

## Discussion

Mapping cell-type-specific 3D genome folding and looping is important for understanding cell-type-specific CRE interactions and gene regulation. To obtain contact maps with sufficiently high resolution to understand CRE looping across cell types, either experimental 3D genome mapping^8,10,11,13,14,16^ or deep learning with existing datasets alone^17–29^ is currently inadequate. In this study, we overcome this challenge and show that integrating deep learning with experiments specifically generated for our designed model architecture is a powerful approach for finescale 3D genome mapping. We developed Cleopatra, an attention-based deep learning model that is pre-trained on experimental genome-wide Micro-C and fine-tuned on selected ultra-high-resolution RCMC, ultimately predicting contact maps at 500bp and 2kb bin sizes. Using Cleopatra, we generate the highest resolution genome-wide contact maps in four human cell types, enabling unprecedented insight into looping and gene regulation.

Cleopatra achieves state-of-the-art performance and comprehensively maps cell-type-specific microcompartments and loops across the human genome. We generate the largest catalog of loops in GM12878, HCT116, K562 and H1 hESCs, identifying over 900,000 loops across the four cell types, approximately half of which are celltype-specific. Although large-scale 3D genome structures such as TADs and A/B-compartments are thought to be conserved across cell types^79^, loops and microcompartments can often be cell-type-specific, and we nominate new TFs that may establish cell-type-specific domain boundaries and loops. Additionally, about half of the loops are anchored on at least one side by “ other” anchors, with “ other” representing anchors that are not annotated as enhancer, promoter or CTCF by ENCODE^31^. The “ other” category is enriched for repressive histone marks, but remains poorly understood, suggesting that our mechanistic understanding of loop formation remains far from complete.

Nevertheless, the approximately quarter million loops per cell type predicted by Cleopatra help to address several open questions in distal gene regulation. First, it remains debated whether enhancers primarily act on the closest gene^56,57^ or frequently skip the closest gene^3–5^. We show that about a third of enhancers do not loop to the closest gene, consistent with estimations from functional genomics data^3^. Second, while developmental enhancers are thought to be more distal and cell-type specific than housekeeping enhancers^64,65^, Cleopatra suggests that housekeeping genes form more long-range E-P interactions that are also frequently cell-type specific. Third, it remains unclear whether distal gene regulation is typically dominated by single enhancers or whether multiple enhancers all contribute to gene expression. We show that promoters form about a dozen loops on average, and that while single enhancers can lead to high gene expression, other loops and enhancers can help to further boost expression levels to a smaller extent. Given that individual E-P loops appear to be infrequent and stochastic^1,80,81^, gene regulation by multiple enhancers is likely to be more robust.

While our approach is the first to allow ultra-deep genome-wide 3D genome mapping, a limitation of our integrated approach is that generating the Micro-C and RCMC training data requires tens of millions of homogeneous cells and Cleopatra requires a significant number of epigenomic tracks. With Micro-C and RCMC data for more cell types, we envision that we can apply Cleopatra to predict RCMC-resolution 3D maps from a limited number of epigenome tracks, ultimately allowing for in-silico prediction of ultra-high-resolution 3D genome structure even from rare cell types and tissues.

## Acknowledgments

We thank Bo Xia, Danilo Dubocanin, Nick Altemose, Geoff Fudenberg, Bin Zhang and Joe Paggi for feedback on the manuscript. We thank all members of the Hansen lab for discussion and support throughout the project, including Jamie Drayton, Jacob Kæstel-Hansen, Miles Huseyin, James Jusuf, Sumin Kim, Masahiro Nagano, Sarah Nemsick, Jack Toppen, and Harvey Yang for feedback on the manuscript. C.K.Y.H is supported by the MIT Novo-Nordisk Artificial Intelligence Postdoctoral Fellowship and National University of Singapore Development Grant. A.S.H. acknowledges funding support from the NIH (DP2GM140938, R33CA257878, R01EB035127, UM1HG011536, R01CA300848, R03OD038390), an NSF CAREER award (2337728), the Gene Regulation Observatory of the Broad Institute of MIT and Harvard, the Novo Nordisk Foundation Center for Genomic Mechanisms of Disease (NNF21SA0072102), a Pew-Stewart Scholar for Cancer Research award, the Mathers Foundation, and an RSC award from the MIT Westaway Fund. J.L. acknowledges funding support from the NIH (R35HG011279 and R03OD038390). This work was supported by the Bridge Project, a partnership between the Koch Institute for Integrative Cancer Research at MIT and the Dana-Farber/Harvard Cancer Center. We thank the MIT Koch Institute’s Robert A. Swanson (1969) Biotechnology Center for technical support, specifically the Integrated Genomics and Bioinformatics Core and MIT BioMicro Center, and this work was supported in part by the Koch Institute Support (core) Grant P30-CA14051 from the National Cancer Institute. We also thank the Walk-Up Sequencing services of the Broad Institute of MIT and Harvard.

## Author contributions

C.K.Y.H, F.F, J.L., and A.S.H. conceived and designed the project. C.K.Y.H performed all experiments and most bioinformatics analyses. V.R. developed CHIRON with input from F.F. and performed sub-pixel localization loop analyses. F.F. developed Cleopatra with input from all authors, and performed benchmarking and eQTL analyses. A.S.H. and J.L. supervised the project. All authors contributed to drafting and editing the paper and figures.

## Data availability

Due to the large amount of raw data generated in this study (11TB), fastqs are being uploaded to SRA and will be available by the time of publication. All processed files for RCMC and Micro-C can be found on Zenodo at https://zenodo.org/records/15303879. Cleopatra predictions (500bp and 2kb) can be found on Zenodo at https://zenodo.org/records/15299993.

## Code availability

Code for processing Micro-C/RCMC data and bioinformatic analyses can be found at https://github.com/ahansenlab/cleopatra analysis code. Code for running Cleopatra can be found at https://github.com/liu-bioinfo-lab/Cleopatra/. Code for CHIRON and associated analyses can be found at https://github.com/ahansenlab/chiron.

