## Supplementary Information for "Genome structure mapping with high-resolution 3D genomics and deep learning"

This document contains:

- Supplementary Notes
- Methods
- Supplementary Figures
- Description of Supplementary Tables

#### Supplementary Notes

##### Supplementary Note 1: Micro-C, RCMC, and accessibility

Because Micro-C and RCMC relies on MNase digestion of chromatin into primarily nucleosomes, it is important to ascertain that Micro-C and RCMC maps report on biologically meaningful looping interactions and not just accessibility artifacts. This has already been validated in prior Micro-C studies[1–6], but here we summarize six key pieces of evidence for validation of Micro-C.

First, all 3D genomics methods including Hi-C and Micro-C show unequal coverage and tend to obtain more ligations in euchromatin than heterochromatin. Such accessibility or coverage bias is easily visible in raw Hi-C and raw Micro-C maps. Importantly, this will introduce a multiplicative bias and multiplicative bias can be corrected by "matrix balancing" typically implemented through the Knight-Ruiz algorithm or ICE[7], which we use here. Thus, unequal coverage is intrinsic to both Hi-C and Micro-C but can be corrected for using ICE.

Second, there is the question of whether ICE works as well on Mb-sized RCMC regions as it does on whole genomes. To test this, we previously compared the effectiveness of ICE matrix balancing on regions vs. whole genomes and found ICE to work as well on RCMC regions (see Extended Data Fig. 6a in Goel *et al.* 2023)[5].

Third, there is the question of accessibility bias specifically for loop calling. If Micro-C and RCMC simply makes loops appear between all accessible regions, then one would expect all ATAC/DNase-peaks to form loop anchors and for all loop anchors to have a DNase peak. However, this is not the case as we show in Figure S3. Specifically, Figure S3 shows three examples: one loop where both anchors are DNase-seq negative and thus not accessible; one where one anchor is DNase-seq negative and thus not accessible and the other is DNase-seq positive; and one where both anchors DNase-seq positive. Similar observations were made in mouse embryonic stem cells, where fewer than half of all ATAC-peaks formed loop anchors and where several loop and microcompartment anchors were ATAC-negative (see Extended Data Fig. 6b-d in Goel *et al.* 2023)[5].

Fourth, if loop anchors were largely determined by accessibility bias in Micro-C but not in Hi-C, then CTCF loops would persist after cohesin depletion in Micro-C but be lost in Hi-C. This is because CTCF remains bound to chromatin after cohesin depletion and thus CTCF sites remain accessible. Prior Hi-C studies have conclusively shown loss of CTCF/cohesin loops without loss of CTCF chromatin binding upon cohesin depletion[8–10]. Exactly the same is seen in Micro-C and RCMC, where cohesin depletion similarly leads to loss of CTCF/cohesin loops without loss of CTCF chromatin binding[4,5,11]. This shows that loops are sensitive to biological perturbations in both Hi-C and Micro-C and that loop anchors are not simply a function of accessibility, because CTCF loop anchor accessibility persists upon cohesin depletion.

Fifth, if accessibility bias was the major determinant of loop formation in Micro-C and RCMC, then one would expect loops to be stronger when anchors are more accessible. However, the opposite is seen in mitosis[6]. RCMC analysis of gene-rich regions during the mitosis-to-G1 transition recently uncovered microcompartments in mitotic chromosomes that are strongly present in prometaphase, peak in strength in ana/telophase, and then weaken as cells enter G1[6]. These microcompartments largely consist of CRE loops anchored by enhancers and promoters. Thus, CRE loops are stronger in ana/telophase than in G1/interphase. However, CRE anchors, enhancers and promoters, are somewhat less accessible in mitosis and more accessible in G1/interphase[12]. Thus, if accessibility was the major determinant of CRE loop strength, then Micro-C and RCMC-observable CRE loops should be stronger in G1/interphase than in ana/telophase, which is opposite to what is experimentally observed[6].

Sixth, even if Hi-C does not have the resolution to detect 3D structural features such microcompartments, it should still be possible to detect microcompartments in Hi-C by genomic averaging. Specifically, microcompartments are thought to form through relatively promiscuous affinity-mediated interactions between CREs through a mechanism that is also called block co-polymer microphase separation in polymer physics[5,6]. If this is true, then averaging the interactions of all CREs in the genome should show above-background signal in Hi-C. Indeed, this is precisely what Friman *et al.* observed through averaging of Hi-C in multiple datasets and under multiple different conditions[13]. Thus, while Hi-C does not have the sensitivity to detect microcompartments individually, the signals do become clearly visible over background when averaged across possible CRE-CRE interactions in the genome as shown in Friman *et al.*[13].

#### Supplementary Note 2: Loop calling and centering in RCMC maps

##### Loop calling with CHIRON

In order to perform accurate analyses of loop distributions in RCMC and provide training information about RCMC loops to Cleopatra, a method that comprehensively calls RCMC loops is essential. In particular, since loops are used to weight Cleopatra's inputs during training (Methods), the full scope of loops present in RCMC should be represented in the loop calls used for model training. However, fine-scale loops and microcompartments are not robustly detected by existing loop callers, necessitating manual annotations in previous RCMC studies[5,6]. To comprehensively call loops in all RCMC regions, we developed CHIRON (**Ch**romatin **I**nteraction **R**ec**o**gnition via **N**eural-nets), a convolutional neural network that is pre-trained on Micro-C loops called by state-of-the-art Micro-C loop caller Mustache[14] and fine-tuned on manually annotated RCMC loops (Figure S5A). We show that CHIRON outperforms Mustache on RCMC maps at 1kb bin size (Figure S5B-S5C), particularly for fine-scale loops and microcompartments. CHIRON achieves a higher area under the precision-recall curve than Mustache for unseen loops in two regions, with the greatest improvement observed for the highest-depth GM12878 loops (Figure S5D-S5E). Thus, CHIRON is a new loop caller that can be applied to accurately identify loops in ultra-high-resolution, dense 3D contact maps.

The RCMC loop set used in this study was comprised of manual annotations from regions 1-5 (which were partially used to train CHIRON) and CHIRON-predicted loops for regions 6-14. These loops were used for all RCMC loop analyses, for benchmarking Cleopatra loop strength against RCMC loop strength, and for weighting Cleopatra's fine-tuning loss function. All loops with annotations can be found in Table S6.

##### Precise loop anchor annotation with fracshift

The precision of loop anchor annotation is typically limited to the bin size at which loops are called. However, loops are not anchored at discrete bin intervals, and we reasoned that the high depth of our maps may allow us to localize the "center" of a loop within a single bin. We applied the fracshift algorithm[15,16] (Figure S6A), commonly used in super-resolution microscopy, to localize loop centers within 200 bp bins (Methods). Briefly, fracshift computes the intensity-weighted centroid of a dot (here, a loop region) over many iterations, shifting the region at each step to center on the centroid coordinate computed in the previous step. This eliminates typical biases associated with centroid computation, which tends to position coordinates towards the edges of pixels. We validated localization precision by showing that fracshift-localized CTCF loop anchors have a smaller median offset from CTCF ChIP-seq peak summits compared to naive loop calls (Figure S6B). However, localization convergence and accuracy decline rapidly upon data downsampling, highlighting that fine-scale loop centering will only be possible with very deeply sequenced data (Figure S6C-S6D).

#### Supplementary Note 3: Benchmarking RCMC against other methods

##### Capture-based methods

Other methods have been developed to specifically enrich enhancers and/or promoters from Hi-C or Micro-C libraries[17–20]. In these methods, Hi-C libraries are subject to capture either by probes targeting regions of interest (e.g. promoter-capture Hi-C) or by antibodies targeting specific regulatory proteins (such as RNA PolII- or CTCF-HiChIP)[20]. These methods have been successfully applied to understand E-P interactions in a variety of contexts. However, a major limitation of capture Hi-C is that rigorous normalization remain challenging. For genome-wide and RCMC maps, normalization relies on standardizing read count to the sum total of each row/column to normalize out differences in digestion, ligation and PCR efficiencies throughout the genome[7,21,22], but this is not possible for these methods. As a result, promoter-capture Hi-C inherently cannot be as accurately normalized, and the resultant loop calls are qualitative rather than quantitative, limiting its utility in quantitative modeling of E-P interactions.

Our benchmarking against other capture-based methods also show that there is wide variation between datasets. In Mifsud *et al.*, promoter capture Hi-C detects a large number of looping interactions relative to RCMC, with a total of 9351 interactions compared to 2595 from the same promoters in RCMC. However, most of the loops that are unique to promoter capture Hi-C do not appear to form loops in RCMC data (Figure S8A), suggesting that the additional loops detected may be false positive interactions. In other datasets, promoter capture Hi-C data from K562 detected only 216 loops in RCMC regions[23], while RNA PolII ChIA-PET detected 1189 interactions[24], relative to the 2866 loops detected by RCMC. Critically, in K562 promoter capture Hi-C did not detect any interactions with *MYC*, while ChIA-PET and HiChIP could not accurately detect *cis*-regulatory interactions (Figure S8C-S8D)[25–27]. Thus, even among the capture-based methods there are very large differences suggesting that these cannot accurately or quantitatively capture all E-P interactions, highlighting the utility for ultra-high-resolution 3D contact maps generated by RCMC.

##### Functional looping interaction analyses

A recent estimate suggests that less than 10% of all looping interactions with *MYC* are functional interactions using H3K27ac HiChIP[27]. However, HiChIP suffers from the same issues as other capture-based methods. Using RCMC loops instead, the percentage of all interactions that have been functionally validated by CRISPRi is 7/24 for K562 and 7/32 for HCT116, with the caveat that several enhancers were validated in a different colon cancer cell line HT29[27], and that not all the enhancers were interacting with *MYC* in HCT116. In K562, functional loops tend to be stronger than other loops, though this does not appear to be true in HCT116 (Figure S8E-8F). Taken together, while loops are a superset of functional interactions, the accuracy of loops in RCMC enables us to get better estimates of the proportion of functional loops, though additional experiments will be needed to generally identify loops with strong transcriptional effects at a given locus.

#### Supplementary Note 4: Rationale for Cleopatra architecture

##### Learning long-distance dependencies between genomic loci

Cleopatra predicts RCMC contact maps at high resolution (500 bp or 2 kb bin sizes), meaning that even contacts between modest genomic distances correspond to long bin intervals - for example, a 250 kb genomic distance spans 500 bins at 500 bp resolution. Accurately predicting interactions between two bins, $i$ and $j$, ideally requires integrating genomic and epigenomic information not only around $i$ and $j$, but also across the entire interval between them.

Traditional convolution-based models, including Akita[28] and DeepC[29], rely on 1D convolutional layers to learn representations for individual bins. However, the receptive field of these convolutions is limited, typically spanning fewer than 20 bins per layer. Expanding the receptive field requires increasing the convolution filter size or stacking many layers, which leads to larger, more difficult-to-train models. As a result, these models primarily capture local signals near bins $i$ and $j$, while neglecting potentially informative features between them, such as insulator elements, which are critical for understanding long-range chromatin interactions.

##### Comparison between Cleopatra and CAESAR

To address this limitation, an effective model must be capable of capturing long-range dependencies. CAESAR[30], for example, uses a graph neural network to achieve this by connecting distant genomic bins based on low-resolution Hi-C data. In this architecture, the Hi-C-derived graph determines which bins are connected. For example, if bins $i$ and $j$ fall within the same topologically associating domain (TAD), they are linked in the graph, allowing the model to propagate information between them. However, this approach depends heavily on the resolution and quality of the Hi-C data, which is far lower than that of RCMC. Replacing Hi-C with Micro-C to build the graph is also infeasible in Cleopatra's setting: since Cleopatra is pre-trained on Micro-C, doing so would result in data leakage.

To overcome these challenges, Cleopatra employs self-attention mechanisms, which allow each bin to attend to all others within the input window (1,250 bins). This design enables Cleopatra to model the full range of dependencies between bins $i$ and $j$, including the intervening regions, without relying on predefined graphs. After passing through convolutional and self-attention layers, the learned representations of bins $i$ and $j$ are concatenated and fed into downstream 2D convolutional layers to predict their interaction.

##### Comparison between Cleopatra and C.Origami

This architecture is conceptually similar to C.Origami[31]. However, Cleopatra differs from C.Origami in several key aspects. First, Cleopatra does not use DNA sequence as input. Despite this, it achieves high predictive accuracy while avoiding unconventional resolutions like 2,048bp ($2^{11}$bp) or 8,192bp ($2^{13}$bp), making its outputs more interpretable and easier to integrate into downstream analyses for biologists. Second, Cleopatra is designed with contact map-centric objectives. To better capture the fine-scale microcompartmental structures characteristic of RCMC, we incorporate insulation scores into the loss function and apply additional weights to loop regions during fine-tuning. Finally, by excluding DNA sequence and adopting a carefully optimized architecture, Cleopatra maintains high performance with a significantly smaller model size than C.Origami. As a result, Cleopatra can generate whole-genome contact maps in under 15 minutes on a standard desktop computer, making it both accessible and practical for widespread use.

### Methods

#### Cell culture

All cell lines were cultured under the same conditions as ENCODE to ensure that the data generated here can be integrated with ENCODE data. GM12878 cells were obtained from Coriell Institute, and K562 cells were obtained from the American Type Culture Collection. Both cell lines were cultured in RPMI 1640 + 15% FBS (HyClone #SH30396.03) + 1% penicillin/streptomycin (Thermo Scientific #1507006) + 2mM GlutaMax Supplement (Gibco #35050061). HCT116 cells were a kind gift from Dr Sarah Johnstone's lab, and were cultured in McCoy's 5A Medium (Sigma-Aldrich #M9309) + 10% FBS + 1% penicillin/streptomycin (Thermo Scientific #1507006). H1 hESCs were obtained from WiCell (also known as WA01, provided by Dr. James Thomson at the University of Wisconsin) and were cultured in mTeSR1 + 1% penicillin/streptomycin (Stemcell Technologies #85850). All cell lines were routinely tested for mycoplasma contamination and found to be negative.

#### Region Capture Micro-C

##### Cell crosslinking

RCMC was performed essentially as described in Goel *et al.* with several modifications[5]. Cells were crosslinked with 3mM Dissucinimidyl glutarate (DSG, ProteoChem c1104-100mg) at a concentration of 1x$10^6$ cells/ml for 35 min at room temperature with gentle mixing. Formaldehyde (Pierce 16% Formaldehyde (w/v), #28908) was then added to a final concentration of 1%, and cells continued to crosslink for 10 min at room temperature with gentle mixing. To quench the reaction, Tris-HCl pH7.5 (Fisher Scientific #503-103-1366) was added to a final concentration of 0.375M with incubation for 5 min at room temperature. Cells were then pelleted for 5 min at 850g at 4°C, and the pellet was washed twice with cold PBS (Thermo Scientific #10010049). Finally, the fixed cells were aliquoted to 5x$10^6$ cells/tube and snap frozen with liquid nitrogen.

##### MNase titration

One 5x$10^6$ pellet from each round of fixation was used for MNase titration. Cell pellets were thawed on ice and resuspended in 500$\mu$l Buffer MB#1 (50mM NaCl, 10mM Tris-HCl pH7.5, 5mM $MgCl_2$, 1mM $CaCl_2$, 0.2% NP-40 Alternative (Millipore Sigma-Aldrich #492018), 1x Protease Inhibitor Cocktail (Sigma-Aldrich #5056489001), then incubated on ice for 20min to extract nuclei. The nuclei were pelleted by centrifugation at 2000g for 5min at 4°C, then washed once with cold Buffer MB#1. The pellet was resuspended with 500$\mu$l Buffer MB#1 and split into 5 tubes of 100$\mu$l each. 4$\mu$l, 7$\mu$l, 12$\mu$l, 20$\mu$l & 30$\mu$l MNase (Worthington Biochemical #LS004798) was added to each tube respectively, then incubated at 37°C for 20 min. Samples were transferred to ice immediately, and MNase was quenched with 4mM EGTA (bioWORLD #40520008) followed by incubation for 10 min at 65°C with shaking. The nuclei were then pelleted by centrifugation at 2000g for 5min at 4°C. To reverse crosslink the DNA, 150$\mu$l of reverse crosslinking solution (1x Tris-EDTA (TE) buffer (Sigma-Aldrich #93183), 1mg/ml Proteinase K (Viagen Biotech #6749722), 10% SDS, 100$\mu$g/ml RNAseA (ThermoFisher #EN0531), 200mM NaCl) was added to each sample, which was then incubated at 65°C for either 2 hours or overnight. Next, the DNA was purified with the Zymo DNA Clean $ Concentrator Kit (Zymo #D4033) according to manufacturer's instructions with one modification - 5 volumes (750$\mu$l) of Binding Buffer was added per tube. Each sample was eluted with 25$\mu$l of Elution Buffer, then quantified by Nanodrop. 1-5$\mu$g of each sample was loaded on a 1.5% agarose gel and each band (monomer, dimer etc) was quantified. The MNase amount that gave ~90% monomer:10% dimer/trimer ratio for each sample was selected.

##### Micro-C

Each RCMC replicate consists of 15-25x$10^6$ cells. For each replicate, Micro-C was performed in tubes of 5x$10^6$ cells each, then pooled just before PCR amplification. All volumes listed below are for 5x$10^6$ cell pellets unless otherwise indicated. All cells/nuclei were pelleted by centrifugation at 2000g for 5 min at 4°C.

To extract nuclei from crosslinked cells, cell pellets were thawed on ice and resuspended in 500$\mu$l Buffer MB#1 and incubated on ice for 20min. The nuclei were pelleted and washed with cold 500$\mu$l MB#1 once, and resuspended in 500$\mu$l MB#1. For MNase digestion, the appropriate amount of MNase (determined by MNase titration) was added, and the samples were incubated for 20 min at 37°C with shaking at 1000rpm on a thermomixer. The samples were transferred to ice immediately and the reaction was quenched with 4mM EGTA and incubation for 10 min at 65°C with shaking. The nuclei were pelleted and washed twice with 1 ml cold MB#2 (50mM NaCl, 10mM Tris-HCl pH 7.5, 10mM $MgCl_2$, 100$\mu$g/ml BSA (Sigma-Aldrich #B8667)).

To generate blunt ends for ligation, nuclei pellets were first incubated in 90$\mu$l of 1x NEBuffer 2.1 (NEB #B7202), 2mM ATP (NEB #P0756), 5mM DTT (Sigma-Aldrich #10197777001) and 25U T4 Polynucleotide Kinase (NEB #M0201) for 15 min at 37°C with shaking. 50U (10$\mu$l) of Klenow Fragment (NEB #M0210) was added and samples were further incubated for 15 min under the same conditions. End labelling was then performed by adding 50$\mu$l of 10x T4 DNA Ligase Buffer (NEB #B0202), 66$\mu$M each of dTTP (Jena Bioscience #NU-1004), dGTP (Jena Bioscience #NU-1003), biotin-dATP (Jena Bioscience #NU-835-BIO14), biotin-dCTP (Jena Bioscience #NU-809-BIOX) and 100$\mu$g/ml BSA (Sigma-Aldrich #B8667) and incubating for 45 min at 25°C with interval mixing (1 min shaking at 1000rpm, 3 min still). The reaction was quenched with 30mM EDTA and incubation at 65°C for 20 min with shaking. Finally, the nuclei were pelleted and washed once with 1ml cold MB#3 (50mM Tris-HCl pH 7.5, 10mM $MgCl_2$ and 100$\mu$g/ml BSA).

Proximity ligation was performed by resuspending nuclei in 500$\mu$l of 1x T4 DNA Ligase Buffer, 100$\mu$g/ml BSA and 10,000U T4 DNA Ligase (NEB #M0202) and allowed to incubate overnight at room temperature with gentle nutation. Nuclei were then pelleted at 3000g for 5 min at 4°C, then resuspended in 1x NEBuffer 1 (NEB #B7001) and 1000U Exonuclease III (NEB #M0206) and incubated at 37°C for 15 min with interval mixing to remove biotin-dNTPs from unligated ends. Reverse crosslinking was then performed by adding 2mg/ml Proteinase K (Viagen Biotech #6749722), 10% SDS (Sigma-Aldrich #L3771), 250mM NaCl and 100$\mu$g/ml RNaseA (ThermoFisher #EN0531) and incubating at 65°C overnight with shaking.

Next, the DNA was purified using the DNA Clean & Concentrator Kit (Zymo #D4033) according to manufacturer's instructions with one modification - 5 volumes (1.59ml) of Binding Buffer was added per tube. The DNA was eluted with 25$\mu$l of Elution Buffer twice (50$\mu$l total), then loaded on a 1% agarose gel. The dinucleosomal bands were extracted using the Zymo Gel Purification kit (Zymo Research #D4008) and eluted in with 25$\mu$l of Elution Buffer twice (50$\mu$l total).

For each 5x10$^6$ cell sample, 50$\mu$l of Dynabeads MyOne Streptavidin T1 (Invitrogen #65601) was washed with 1ml TBW buffer (1M NaCl, 5mM Tris-HCl pH 7.5, 500$\mu$M EDTA, 0.1% Tween-20), then resuspended in 150$\mu$l 2X BW buffer (2M NaCl, 10mM Tris-HCl pH7.5, 1mM EDTA). The beads were mixed with eluted DNA from the gel extraction and water was added to a final volume of 300$\mu$l. The beads were nutated overnight at room temperature to allow binding of biotinylated DNA.

After DNA binding was complete, the beads were washed twice with 950$\mu$l TBW buffer and once with 10mM Tris-HCl pH7.5. Illumina library preparation was performed on bead-bound DNA with the NEBNext Ultra II Kit (NEB #E7645) according to the manual, except incubations were performed with interval mixing (1 min shaking at 1000rpm, 3 min still). Finally, the beads were washed twice with 950$\mu$l TBW buffer and once with 10mM Tris-HCl pH7.5, and resuspended in 115$\mu$l of elution buffer (10mM Tris pH 8.5, 0.1mM EDTA).

To determine the minimum number of PCR cycles required to amplify the library for probe capture, a test PCR was performed with 1$\mu$l bead-bound DNA with the KAPA HiFi HotStart ReadyMix enzyme (Roche #07958927001) and quantified on an agarose gel. All beads were then separated into 6 PCR reactions and amplified for 7-9 cycles with unique index primers (NEB #E7335, #E7500, # E7710) for sequencing. The PCRs were pooled and streptavidin beads were removed before PCR purification using 0.9x Ampure XP beads (Beckman Coulter #A63880). Libraries were quantified with qPCR using the KAPA SYBR FAST qPCR Master Mix (2X) Universal (#KK4601) and Library Quantification Kit standards (#KK4903).

In total, we performed 4 replicates of Micro-C each for GM12878, HCT116 and K562 respectively. For RCMC, we performed 40 replicates for GM12878, 10 replicates for HCT116, 4 replicates for K562 and 4 replicates for H1. Note that not all regions were captured for all replicates. Detailed information about the number of replicates for each region can be found in Table S1.

##### Region selection

Regions for RCMC were selected by manual inspection of available Micro-C or Intact MNase Hi-C (ENCODE ENCSR916MFV & ENCSR477GZK). The first five regions (region1-region5, Figure S1B, Table S1) were selected for to be maximally distinct from each other and include highly active, gene-dense regions and repressed, gene-poor regions. The rest of the regions include regions where CRISPRi data have been generated (region6, *MYC* gene) and regions that appear visually very different between cell types.

##### Epigenome annotations and ChromHMM analysis

To validate the diversity of input regions, we split the genome into 5kb bins and created a vector of 12 epigenomic tracks (DNase-seq, CTCF, RAD21, H3K4me1, H3K4me2, H3K4me3, H3K9ac, H3K9me3, H3K27ac, H3K27me3, H3K36me3, H3K79me2). The bins were then embedded into 2-dimensional space with tSNE and compared with chromHMM annotations. chromHMM annotations were downloaded from ENCODE (Table S4). Details of all regions can be found in Table S1.

##### Library capture & sequencing

For library capture, 80-mer biotinylated probes tiling the selected regions was purchased from Twist Biosciences. Three sets of probes were purchased for the 14 regions. The probe sets cover regions1-2, regions 3-5, and regions 6-14 respectively. All coordinates covered by the probe sets can be found in Table S3. Capture was then performed according to Twist Bioscience's Standard Hybridization Target Enrichment Protocol, except that a test PCR was performed with 1.5$\mu$l of DNA before final PCRs to determine the minimum number of PCR cycles required. The entire sample was split into 2 PCR reactions and amplified for 3-5 reactions, then purified according to the Twist protocol. Finally, libraries were quantified by qPCR as described above. Libraries were sequenced on the Illumina NovaSeq X 10B or 25B, 2x150 paired-end reads, at the Broad Institute of MIT and Harvard's Walk-Up Sequencing.

##### Data processing

Paired-end reads were aligned with bwa-mem2 (v2.2.1) with the `-SP` flag to the UCSC hg38 AnalysisSet genome and processed with pairtools (v1.0.2) `parse2` (flags: `--add-columns mapq --expand --report-position outer --min-mapq 30 --max-insert-size 150 --drop-sam`). The resulting pairs files were filtered to keep only reads with both ends within the RCMC regions using a custom script (filter_reads_merged.py). Reads from the same replicate (from different regions) were merged, then deduplicated with pairtools `dedup` (flags: `--max-mismatch 1`). Files from the same replicates were then merged, indexed with pairix (v0.3.7), and made into a .cool file with cooler (v0.10.2) `cload pairs` at 50bp bin size. Finally, .cool files were converted to .mcool files using cooler `zoomify` with the `--balance` flag up to 10Mb bin sizes. For Micro-C data, all the steps were the same except the step where reads are filtered for RCMC regions.

To calculate the resolution of our contact maps using the definition provided in Rao, Huntley *et al.* 2014, contact maps were binned at 50, 100, 150, 200, 250, 300, 400, 500, 800, 1000bp and unbalanced (raw) matrices were obtained using cooltools. We then calculated rowsums (corresponding to reads in each bin), and calculated the proportion of bins that have reads >1000 reads for each resolution and region. The resolution reported for each region in Table S1 corresponds to lowest bin size where the proportion of bins with >1000 reads is >80%.

#### Cleopatra model details

##### Micro-C and RCMC data pre-processing

To avoid having the model's loss function dominated by short-range interactions, Micro-C and RCMC contact maps were first converted to observed/expected (OE) maps. The average values of each contact distance (at 500bp and 2Kb resolutions) after ICE normalization were stored as an "expected vectors". Since the average contact at longer distances were too small and dividing by the small average values will result in very large OE values, we set a threshold of $10^{-5}$ so that all expected vector values smaller than $10^{-5}$ were replaced with $10^{-5}$. All contact values were then divided by the corresponding expected vector values for a given distance. We then calculated $log10(x + 1)$ for each pixel to reduce magnitude differences. To overcome the sparsity of contact maps at 500bp resolution, we smoothed the contact map with a 5×5 uniform kernel. Finally, all values greater than 6 are replaced with 6.

##### Model inputs and pre-processing

We selected 19 common input features that were availble for all four cell types: DNase-seq, H3K4me1, H3K4me2, H3K4me3, H3K9ac, H3K9me3, H3K27ac, H3K27me3, H3K36me3, H3K79me2, H4K20me1, CTCF, EZH2, POLR2A, JUND, REST, RAD21, H2AFZ, and phastCons score The *.BigWig* files were downloaded from ENCODE according to Table S5, which were all processed with the ENCODE4 pipeline in hg38, and the epigenomic signals were binned at 500bp and 2kb bin sizes with the Python package *pyBigWig* and stored as *numpy* arrays. PhastCons data was also downloaded as *.BigWig* files and binned with the same approach. All signals were normalized using GM12878 as a reference, i.e., for each epigenomic signal, the arrays of other cell types (e.g., K562) were multiplied by a constant to ensure the vector's average values were the same as GM12878. We calculated $log10(x + 1)$ for each value to reduce magnitude differences, and replaced all values greater than 6 with 6.

In each mini-step, the model takes a 1,250-bin region (625kb for 500 bp resolution, 2.5Mb for 2 Kb resolution). This input is a 19×1,250 matrix, where 19 is the number of genomic features used. The same positional encoding as the Transformer model was used[32], which is an 8×1,250 matrix. To account for interaction distance, we also added distance encoding as part of the inputs, which is a 1,250×1,250 matrix where the value at the $x$-th row and $y$-th column is $|x - y|$.

##### Model structure

The genomic features and positional encoding matrices are concatenated to form a (19+8)×1,250 matrix (Input1D matrix), which corresponds to 625kb for 500bp Cleopatra and 2.5Mb for 2kb Cleopatra in genomic distance. The distance embedding matrix is passed through an embedding layer, which converts the numeric distances (e.g., 0, 1, 2, ..., 1,250) to 8-dimensional embeddings. This 1,250×1,250×8 tensor is referred to as the DistanceEmb tensor. The Input1D matrix is passed through three convolution blocks (Conv1D layer + BatchNormalization layer) with 64 convolution kernels with a size of 21. The output shape of each group is 64×1,250. The output is then passed through two attention blocks (MultiHeadAttention layer + BatchNormalization layer), with 8 attention heads and 64 total output dimensions. The output shape of each group is 64×1,250. All convolutional layers use the *relu* activation function. The outputs of all previous layers are concatenated into a 320×1,250 matrix (i.e., a 320-dimension representation for each bin). For each pair of bins $i$ and $j$, the representations of bin $i$ and bin $j$ are concatenated to form a vector, which passes through a dense layer with a dimension of 24, resulting in a 1,250×1,250×24 tensor. This tensor is concatenated with the DistanceEmb tensor to form a 1,250×1,250×32 tensor, which then passes through two Conv2D blocks (Conv2D layer + BatchNormalization layer), with 24 kernels and 3×3 kernel sizes, resulting in a 1,250×1,250×24 tensor. The output then goes through the final layer to convert the 24-dimensional embedding into the predicted OE contact value of each pixel.

##### Cleopatra model training and fine-tuning

For model training, the genome was segmented into windows of 1,250 bins (625kb for 500bp bin size and 2.5Mb for 2kb bin size), with each window serving as an individual training example. Micro-C data for all chromosomes was used for pre-training, and the mean squared error (MSE) loss function was used with a learning rate of $10^{-3}$.

For fine-tuning, we first used 10 regions (regions1, 2, 5, 7-10, 12-14) and held out 3 regions (regions4, 6, 11) to evaluate the accuracy of the predictions. Region2 was not used for training or fine-tuning because the data quality was much lower than the others (see Table S1). Additional weights (i.e., penalty in loss functions) for all RCMC loops were added by doubling the loss function for 11×11 bin squares around loops. We found that in addition to weighted MSE, insulation loss (defined as the MSE of predicted and RCMC diamond insulation scores) was helpful in generating more experimentally realistic maps. Diamond insulation scores were calculated at 500bp and 2kb resolutions for 500bp and 2kb Cleopatra respectively, with a window size of 10 bins. We tested three strategies in fine-tuning: (1) freezing the 1D convolutional and self-attention layers, and re-initializing the last output 2D convolutional layers, (2) keeping the weights of all original model layers, and only retraining the last output 2D convolutional layers, and (3) keeping the weights of all original model layers, and unfreezing all weights during fine-tuning. We found that the third strategy resulted in the best model performance. A learning rate of $5 \times 10^{-5}$ was used in fine-tuning. Once we validated the accuracy of Cleopatra, we then retrained Cleopatra using all regions (except region2) to obtain a final model, and applied it to generate the genome-wide predictions. During prediction, Cleopatra outputs contact maps of 1,250-bin sliding window with a 250-bin step length, which effectively covers interactions within 1,000 bins (500kb for 500bp bin size and 2Mb for 2kb bin size). The output matrices were merged to generate continuous predictions across the genome by averaging across pixel values corresponding to the same genomic bins.

##### Converting Cleopatra predictions to .cool files

The outputs of Cleopatra predictions are log(OE) values over 500kb or 2Mb sliding windows. To generate contact maps that resemble experimental maps the values were converted to "observed" values by multiplying each value by the expected value of that bin. For the holdout regions, the expected vectors were calculated from the respective RCMC regions. For the genome-wide predictions, we first fitted a curve with RCMC-available regions, which uses a linear function to estimate RCMC expected values of each contact distance from Micro-C expected values and DNase-seq signal. Then, the RCMC expected values for each 100kb regions was estimated with the curve for the entire genome, and used to multiply by the predicted OE values. For the genome-wide predictions, since true OE vectors are not available for regions without RCMC, we fitted a linear function to estimate RCMC expected values of each contact distance for each 100kb region. This function was fitted with data from RCMC-available regions, and takes Micro-C expected values and DNase-seq signal as inputs.

The outputs of the holdout region models were merged into a single contact map to facilitate analyses using existing tools. Each matrix of "observed" values was first converted into a .cool file with the appropriate genomic coordinates. The .cool files were then merged into one file using cooler `merge` with `agg=mean` to average values that were predicted in multiple matrices. Genome-wide predictions were already concatenated as described above, so .cool files with "observed" values for each chromosome were first generated, then merged into one file with cooler `merge`. Finally, .cool files were converted to .mcool files using cooler `zoomify` up to 10Mb bin sizes.

##### Comparing Cleopatra maps to RCMC

To evaluate the similarity between RCMC and Cleopatra matrices, Pearson's correlations were calculated for each distance from the diagonal at 500bp or 2kb bin sizes for 500bp and 2kb Cleopatra respectively. The correlations were then plotted with the `geom_smooth` function in R using the ggplot2 package[33].

Loop strength calculation was performed by summing the values obtained from `ObsExpSnipper` in cooltools (v0.7.0) using loops identified from RCMC maps. Loop strength was calculated using the respective model bin sizes (500bp or 2kb) with 3kb and 5kb loop anchors respectively in both RCMC and Cleopatra maps.

##### Loop pileups on Cleopatra models with fewer inputs

Pileups were performed with loops identified from RCMC maps. To identify loops anchored by CTCF, H3K4me3 and H3K4me1, loops were overlapped with peak calls from the respective ChIP-seq datasets (Table S5), and only loops where both anchors overlapped a ChIP peak was kept. Loop pileups were then performed using coolpuppy (v1.1.0)[34] with a window size of 20kb around the midpoint of each anchor.

##### Comparisons with other models

The trained Akita v2 model[28,35] was downloaded from GitHub and the tutorial notebook was used to obtain Akita predictions. The holdout region6 in Cleopatra overlaps with some of the Akita test regions, which we used for comparison with the Cleopatra holdout model (chr8:127410000-128458576).

C.Origami predictions were obtained using the provided Google Colab tutorial notebook for 2 test regions[31] (chr10:122700000-124797152, chr15:59100000-61197152). Since these regions do not overlap RCMC holdout regions we compared Cleopatra with C.Origami using Hi-C and Micro-C data.

#### Data analysis

##### Data visualization and plotting

All 1D tracks and 3D contact maps were visualized in Python 3.9 or R v4.4.1 using a modified version of plotgardener[36]. To visualize data from .mcool files, data was dumped from regions of interest using the cooler `dump` function with the flags `--join --balanced --header --range <coords>`. For Cleopatra maps the `--balanced` flag was not used.

##### Comparing with published Micro-C/Hi-C Data

Sources of other 3D genomics datasets used for comparisons can be found in Table S4. For H1 Micro-C[3] and GM12878 Hi-C[37] data, .pairs files were downloaded and processed using the same steps as above (see Data Processing). For HCT116, K562 and H1 Hi-C data, .mcool files were downloaded from the respective sources and used as is.

To calculate distance decay from the diagonal, the cooltools `expected_cis` function was used with the flags `smooth=True, aggregate_smoothed=True, smooth_sigma=0.1` at 1kb bin size[38]. Curves were truncated at 3x$10^6$bp because the capture regions are around 2-3Mb in size. To accurately compare datasets contact probability for Micro-C/Hi-C was only calculated only within the regions selected for RCMC.

##### Boundary calling and analysis

Boundaries were called with cooltools (v0.7.0) with the `insulation` function at 200bp bin sizes using a window size of 20kb, followed by filtering for `is_boundary == TRUE`. In RCMC data, false positive boundary calls sometimes occur where there are visible gaps in the data (due to lack of probe coverage in those gaps), thus, boundaries overlapping these gaps were further filtered.

For the boundary comparison across cell types, the ComplexHeatmap package (v2.21.0)[39,40] was used to to identify unique/shared boundaries and to generate the UpSet plot. For motif analyses, AME[41] was used to calculate the enrichment of motifs relative to random 200bp sequences that are not within 10kb of a boundary in any cell type. The motifs used were downloaded from HOCOMOCO v12 Core (H12CORE_MEME_format.meme)[42]. While representative motifs were plotted, additional motifs can be found in Table S2. The enrichment of ChIP signals at RCMC/Micro-C boundaries was plotted with EnrichedHeatmap (v1.34.0) using scores from the respective bigWig files downloaded from ENCODE with `mean_mode='w0'`.

##### Downsampling RCMC

Only GM12878 RCMC was downsampled because it is the most deeply sequenced dataset. Downsampling was performed with pairtools `sample` at the indicated fractions from the final RCMC .pairs file.

##### Loop calling with CHIRON

RCMC loops were identified using the **Ch**romatin **I**nteraction **R**ec**o**gnition via **N**eural-Net (CHIRON) loop caller, a convolutional neural network built to classify loops in RCMC data. CHIRON predicts the probability of a loop existing at each pixel of an image representation of a genomic contact map. The model takes in 31x31 pixel regions of a contact map binned at 1kb. Contact maps are observed/expected normalized and each loop region is max-normalized to ensure signal ranges are consistent for each data point. Loop regions are then passed through three convolutional layers and two dense layers with *relu* and *softmax* activation functions. The output is a list of high-probability loop coordinates. A single loop center is then selected from each region of contiguous high-probability pixels via local maximum finding in the contact map.

CHIRON was pre-trained on 71,776 Micro-C loops called by Mustache and fine-tuned on 8,138 manually annotated loops from this study and other RCMC work from this lab. Manual annotation for this study was performed using a custom-written graphical user interface, while prior studies utilized Hi-Glass. The Micro-C data included HEK293 cells from Narducci and Hansen[43], mESCs from Jusuf *et al.*[44], and H1 hESCs from Krietenstein *et al.*[3]. The RCMC included 3,376 loops from region1, region2, and region4 from all four cell types in this study, 1,226 loops from Goel *et al.* in mESCs[5] and 3,211 loops from mouse erythroblasts[6], and 325 loops from human erythroid cells (unpublished). All loops (positive training data) were matched with a non-loop region at the same interaction distance (negative training data). For fine-tuning, the weights of all layers until the last convolutional layer were frozen such that CHIRON retained basic loop features learned from Micro-C and only higher-order RCMC-specific features were learned from RCMC.

Manual annotations from a loop-rich and a loop-poor locus were held out for model evaluation: region3 (696 loops across cell types) and region5 (14 loops across cell types). CHIRON was benchmarked against Mustache by comparing each method to manual annotations at 500bp, 1kb, and 2kb binsizes in the held-out regions. A 3kb Euclidean distance cutoff between a called loop coordinate and a ground truth loop coordinate was considered a true positive. Precision-recall curves were generated for CHIRON across probability cutoffs and for Mustache across FDR cutoffs. The specific Mustache command was: `mustache -r 1kb -pt 0.1 -sz 1.6 -st 0.88`.

##### Loop center localization

200bp loop anchors were computed using the fracshift algorithm[15,16] with some modifications. For each loop, a base-pair center at 200 bp binsize was computed as follows. First, an initial center was defined as the local maximum of the 1-pixel region around the annotated center of the loop at 800 bp binsize. A fractionally centered coordinate was computed as the intensity-weighted centroid of a circular mask around the loop center with a 3-pixel radius at 200 bp binsize, repeated for 20 iterations. Each iteration computes the centroid of the region centered at the fractional center coordinate from the previous iteration. Finally, the coordinate was converted from image pixels back to base pair coordinates.

To benchmark the centers obtained by fracshift against standard loop calling methods, we assessed the positioning of CTCF-CTCF loop anchors relative to CTCF ChIP-Seq peak summits. We filtered loops for those that had CTCF motifs and a CTCF ChIP peak within a 2kb region around the loop center. In particular, we used the 'known1' CTCF motif set from Kheradpour *et al.*[45] (converted from hg19 to hg38 using UCSC liftOver) and the CTCF ChIP peak summits from ENCODE ENCSR000DZN (narrowPeak .bed file). Loop centers for manual annotations and fracshifted loops were the basepair coordinates obtained from the annotator GUI described above or the fracshift algorithm, respectively. As a benchmark, loop centers for Mustache were the centers of the 1kb bins obtained by calling Mustache with default parameters. For comparisons in all 14 regions, only loops that agreed between Mustache and CHIRON (Mustache loops within 3kb Euclidean distance from a CHIRON loop and vice versa) were used for analysis to eliminate potential false positives in either set. For comparisons in the first 5 regions, all CTCF-annotated loops were evaluated as there were relatively fewer loops to analyze.

To evaluate the performance of fracshift at different read depths, we used the GM12878 downsampled data described above. We restricted analyses to region1 because read depths differ with different regions, so absolute read numbers across downsampled datasets can only be evaluated for one region with respect to itself. We defined convergence for a given loop if the loop center deviated by less than 0.1 pixels for the 3 final fracshift iterations. This metric was chosen based on the performance of fracshift on simulated dots, for which the deviation of computed vs. true centers was around 0.1 pixels. Distance of downsampled centers from original centers was computed via Euclidean distance. Read counts for the GM12878 region1 in RCMC and Micro-C were computed using `filter_reads_merged.py` as described above.

##### Loop analyses

Loop pileups were performed using coolpuppy (v1.1.0)[34] with a window size of 20kb around the midpoint of each anchor.

All loop analyses use loops at 1kb bin size (1kb flanking the midpoint of the loop call) unless otherwise specified. To classify loop anchors, candidate *cis*-regulatory element (cCRE) annotations from SCREEN v3 (Search Candidate *cis*-Regulatory Elements by ENCODE)[46] were downloaded for each cell type. cCREs with "Promoter-like signatures" (PLS) were considered promoters, cCREs with proximal or distal "Enhancer-like signatures" (pELS, dELS) were considered enhancers, and cCREs with "CTCF-bound" annotations were considered CTCF sites. Since each 1kb anchor may overlap more than one cCRE (which can be smaller than 1kb), anchors were overlapped with all cCREs using InteractionSet R package (v1.33.0)[47] and annotated in the following order: promoter, enhancer, CTCF. For example, if an anchor overlaps both promoter and enhancer annotations, it will be annotated "promoter". Anchors that do not overlap any of these annotations were considered "other".

Loops are considered shared across cell types if both anchors overlap by at least 1.5kb using an anchor size of 5kb. A 4-way comparison was performed using a custom script based on bedtools `pairtopair`[48].

Histone modification scores for each 1kb anchor was calculated using bedtools `multicov` with the respective bam files for each cell type. The scores were averaged across loop classes and z-scored for plotting using the ComplexHeatmap package (v2.21.0).

Loop strength calculation was performed by summing the values from `ObsExpSnipper` in cooltools (v0.7.0). For RCMC, loop strength was calculated using 1kb bin sizes with 3kb loop anchors.

For gene expression analyses, total RNA-seq data was downloaded for the respective cells lines from ENCODE (Table S4). Expressed genes are defined as having FPKM >1.

For the enhancer skipping analyses, enhancers were defined as cCREs with proximal or distal "Enhancer-like signatures" (pELS, dELS) from SCREEN v3 annotations (described above). Enhancers were considered as "skipping" the nearest gene if it does not loop to both of its closest promoters on either side and forms a loop with a gene that is further away. Genes that were less than 5kb to the enhancers were excluded from the analyses.

To identify motif pairs enriched in cell-type-specific E-P pairs, all motifs present in all E-P loop anchors at 1kb were first found using FIMO[49] with the HOCOMOCO v12 Core (H12CORE_MEME_format.meme) database[42]. For each E-P loop, all motifs on one anchor were paired with all motifs in the other anchor, and the co-occurrence of each pair was calculated for cell-type-specific and shared loops. For each motif pair, a chi-squared test was performed to calculate the enrichment of motif pairs over the occurrence of each individual motif separately. Only motif pairs that were significantly enriched over both motifs after Benjamini & Hochberg correction (adjusted p-value <0.05) were kept.

##### Promoter capture Hi-C comparisons

Promoter capture Hi-C data from Mifsud *et al.*[17], Li *et al.*[24] & Caragine *et al.*[27] was downloaded and lifted over from hg19 to hg38 coordinates using UCSC liftover. For details of the datasets see Table S4. To identify loops shared between RCMC and other datasets, bedtools `pairtopair` was used with the parameters `-f 0.5`. For the loop strength comparison with Mifsud *et al.*, the significant interactions file was downloaded and the log(observed/expected) score was used.

##### Loop calling with Mustache

For Micro-C maps, loops were called using Mustache[14] at 1kb bin size with default parameters.

To apply Mustache to Cleopatra maps the mustache script was modified to allow loading of Cooler files without the `balance==False` flag. Loops were then called[14] at 500bp and 2kb for the 500bp and 2kb models respectively, with the flags `--st 0.9 --pt 0.002 --fdr 0.01`. Visually, false positive calls often happen where 'empty' stripes can occur in RCMC or Micro-C data due there are alignment issues arising from differences between the cell lines and the reference genome. To remove these false positives, loops were further filtered to remove all loops within 4 bins of 'empty' stripes in RCMC (for loop calling within RCMC holdout regions) or 8 bins in Micro-C data (for which typically occur). Finally, loops from 500kb and 2Mb predictions were merged using the 500kb predictions for loops <495kb and 2Mb predictions for loops from 495kb to 198kb. All Cleopatra loops with annotations can be found in Table S7.

##### CRISPRi comparisons

For K562 data CRISPRi data comparisons, the processed, harmonized and annotated (Regulated/not regulated) data from Gschwind *et al.* was used because consistent and rigorous thresholds were applied[50]. Loop pileups were performed with coolpuppy[34].

##### Contact enrichment analysis for GM12878 eQTL

The eQTL data was downloaded from GTEx V8[51]. Since most eQTLs are related to linkage disequilibrium instead of regulatory elements, the list was filtered such that for each gene, only the eQTL with the smallest p-values in a $\pm$ 25kb window was kept, leaving 7,690 eQTL-gene pairs. The predicted OE contact values for the filtered interactions was extracted from all four cell types for plotting.

### Supplementary Figures

A

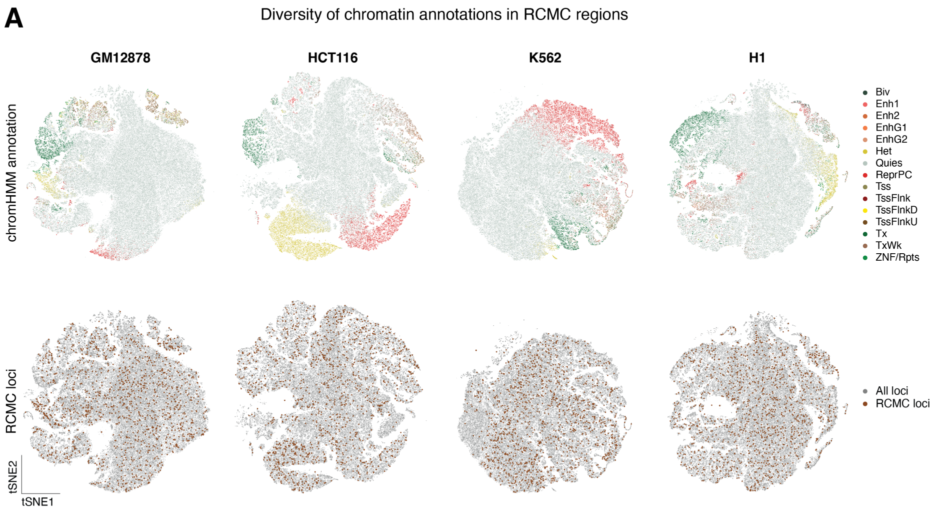


B

Regions selected for RCMC

| | Coordinates (hg38) | Size (Mb) | Number of genes |
|---|---|---|---|
| region1 | chr6:25111952-28621868 | 3.59 | 186 |
| region2 | chr19:36027639-39756265 | 3.73 | 180 |
| region3 | chr5:157036231-160147469 | 3.11 | 62 |
| region4 | chr1:207626481-210338298 | 2.71 | 48 |
| region5 | chr4:61369236-64434279 | 3.07 | 16 |
| region6 | chr8:124988367-129795630 | 4.81 | 55 |
| region7 | chr6:29678698-32256197 | 2.58 | 239 |
| region8 | chrX:47081617-49442070 | 2.36 | 127 |
| region9 | chr1:237286814-240531042 | 3.24 | 35 |
| region10 | chr7:10480243-13506262 | 3.03 | 27 |
| region11 | chr8:62918949-66566105 | 3.65 | 54 |
| region12 | chr4:181289951-184014354 | 2.72 | 39 |
| region13 | chr3:118624552-121533938 | 2.91 | 58 |
| region14 | chr9:106603383-109909256 | 3.31 | 50 |
| **total** | | **44.82** | **1176** |

**Supplementary Figure 1: RCMC regions represent diverse epigenomic features.** (A) t-SNE plot of the human genome's epigenomic landscape for each cell type. Twelve epigenomic tracks (DNase-seq, CTCF, RAD21, H3K4me1, H3K4me2, H3K4me3, H3K9ac, H3K9me3, H3K27ac, H3K27me3, H3K36me3, H3K79me2) from the entire genome are used for the tSNE embedding. Each point represents a 5kb region. The points are colored by ChromHMM annotations[52] (top) or whether the region is part of a selected RCMC region (bottom). (B) Overview of regions selected for RCMC. More details can be found in Table S1.

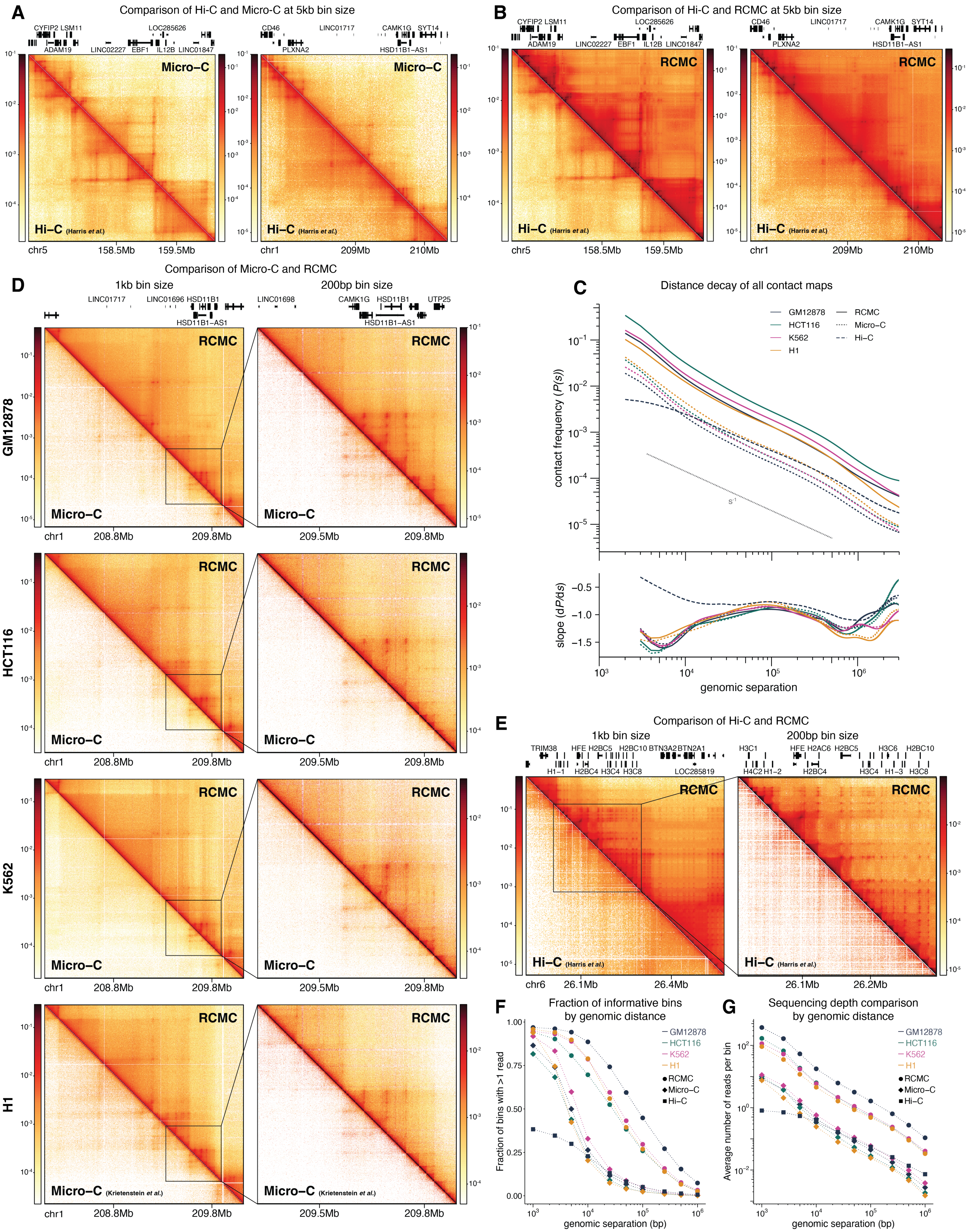

A
Comparison of Hi-C and Micro-C at 5kb bin size
Micro-C
Hi-C (Harris et al.)
chr5
158.5Mb
159.5Mb
chr1
209Mb
210Mb
B
Comparison of Hi-C and RCMC at 5kb bin size
RCMC
C
Distance decay of all contact maps
GM12878
HCT116
K562
H1
RCMC
Micro-C
Hi-C
contact frequency (P(s))
slope (dP/ds)
genomic separation
D
Comparison of Micro-C and RCMC
1kb bin size
200bp bin size
GM12878
HCT116
K562
H1
208.8Mb
209.8Mb
209.5Mb
Micro-C (Krietenstein et al.)
E
Comparison of Hi-C and RCMC
chr6
26.1Mb
26.4Mb
26.2Mb
F
Fraction of informative bins by genomic distance
Fraction of bins with >1 read
genomic separation (bp)
G
Sequencing depth comparison by genomic distance
Average number of reads per bin

**Supplementary Figure 2:** (Figure on previous page) **RCMC outperforms current best Hi-C and Micro-C maps.** (A) Comparison of Hi-C and Micro-C maps at 5kb bin size in GM12878. (B) Comparison of Hi-C and RCMC maps at 5kb bin size in GM12878. (C) Distance decay and derivatives of indicated contact maps in RCMC regions. (D) Representative region comparing RCMC to best available Micro-C data in four cell types. (E) Representative region comparing RCMC to best available Hi-C (33B unique pairwise interactions[37]) in GM12878. (F) Fraction of 200bp bins filled by distance in RCMC, Micro-C and Hi-C (GM12878). (G) Average number of reads per 200bp bin by distance in RCMC, Micro-C and Hi-C (GM12878). All GM12878 Hi-C and H1 Micro-C data used in this figure was obtained from Harris *et al.*[37] and Krietenstein *et al.*[3] respectively.

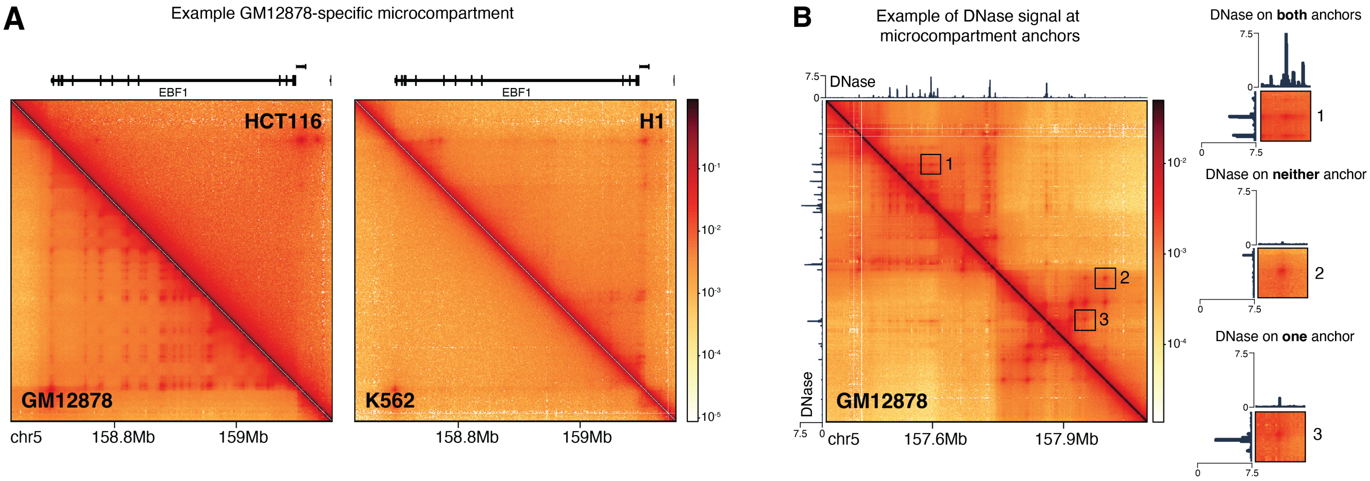


**Supplementary Figure 3: Microcompartment anchors are cell-type-specific and not an artifact of open chromatin regions.** (A) Example cell-type-specific microcompartment in GM12878. Contact maps are plotted at 1kb bin sizes. (B) Representative region showing three loops with varying DNase signal at each anchor. Each loop is an inset corresponding to the indicated loop on the map.

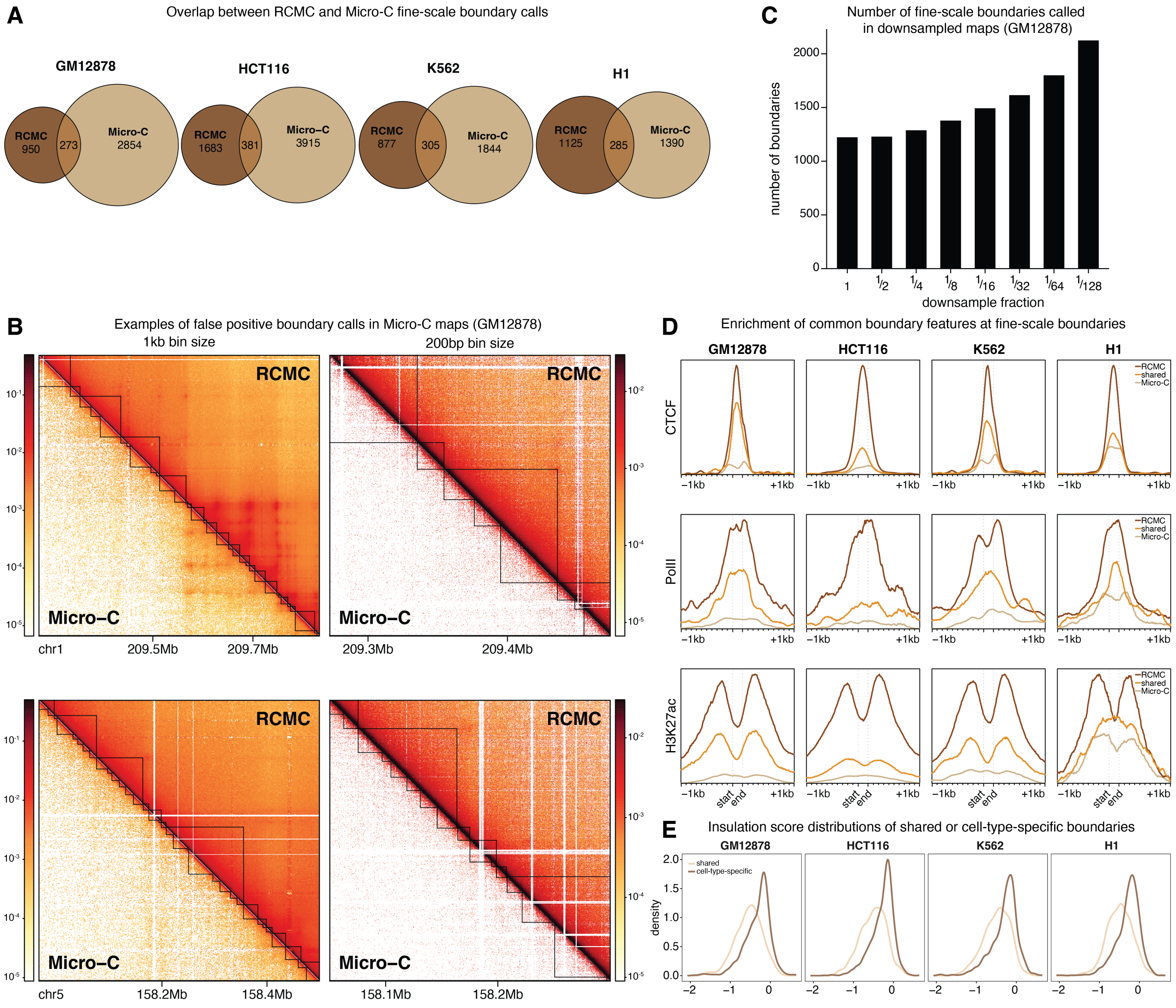


**Supplementary Figure 4: RCMC accurately detects compartment boundaries.** (A) Number of compartment boundaries identified in RCMC, Micro-C (in RCMC regions only) or shared in each cell type. (B) Two representative examples of false positive compartment calls in Micro-C compared to RCMC. The 200bp plots are insets of the 1kb plot. (C) Number of boundaries called in RCMC regions from downsampled contact maps. (D) Metaplot of CTCF, PolII or H3K27ac ChIP-seq for RCMC-specific, Micro-C specific or shared boundaries. Start and end represent the ends of the boundaries, and plots are extended 1kb on either side. (E) Distribution of insulation scores of shared or cell-type-specific boundaries. Lower $\log_2$ insulation scores represent stronger boundaries and vice versa[38].

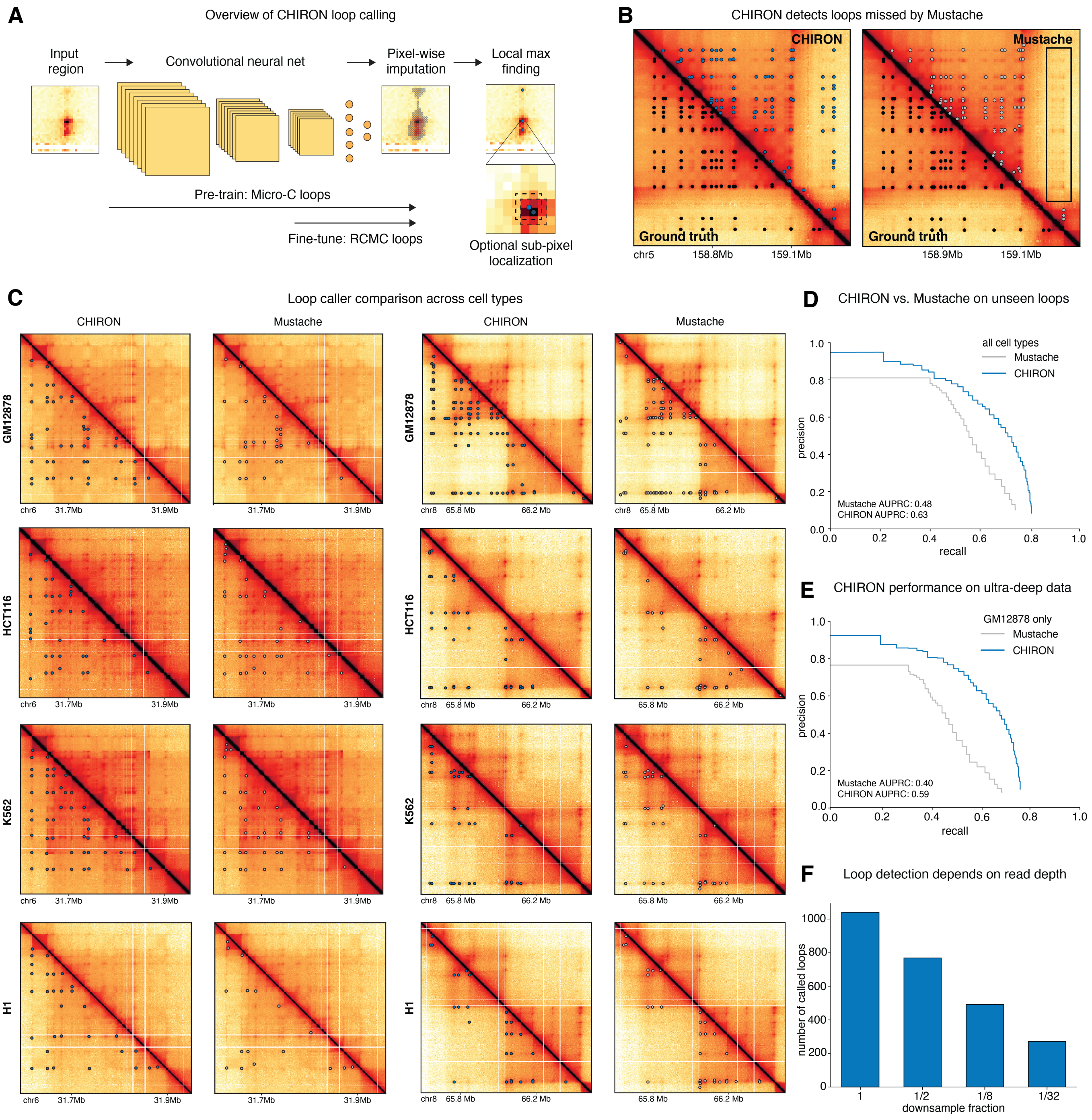


**Supplementary Figure 5: CHIRON outperforms Mustache on RCMC maps.** (A) Overview of CHIRON method for loop calling in highly dense 3D contact maps, including an optional step for loop centering by sub-pixel localization. (B) CHIRON outperforms state-of-the-art Mustache loop caller on a holdout RCMC region. Contact maps are plotted at 1kb bin size. Ground truth, CHIRON and Mustache loop calls are indicated by black, blue and gray circles respectively. (C) Two representative examples of loop calls by CHIRON or Mustache on the same RCMC data in all cell types. Contact maps are plotted at 2kb binsize. (D)-(E) Precision-recall curves for Mustache and CHIRON on the validation set for all celltypes (D) or GM12878 (E). The area under the curve (AUPRC) is indicated. (F) Number of loops called by CHIRON in GM12878 data (region3) after downsampling.

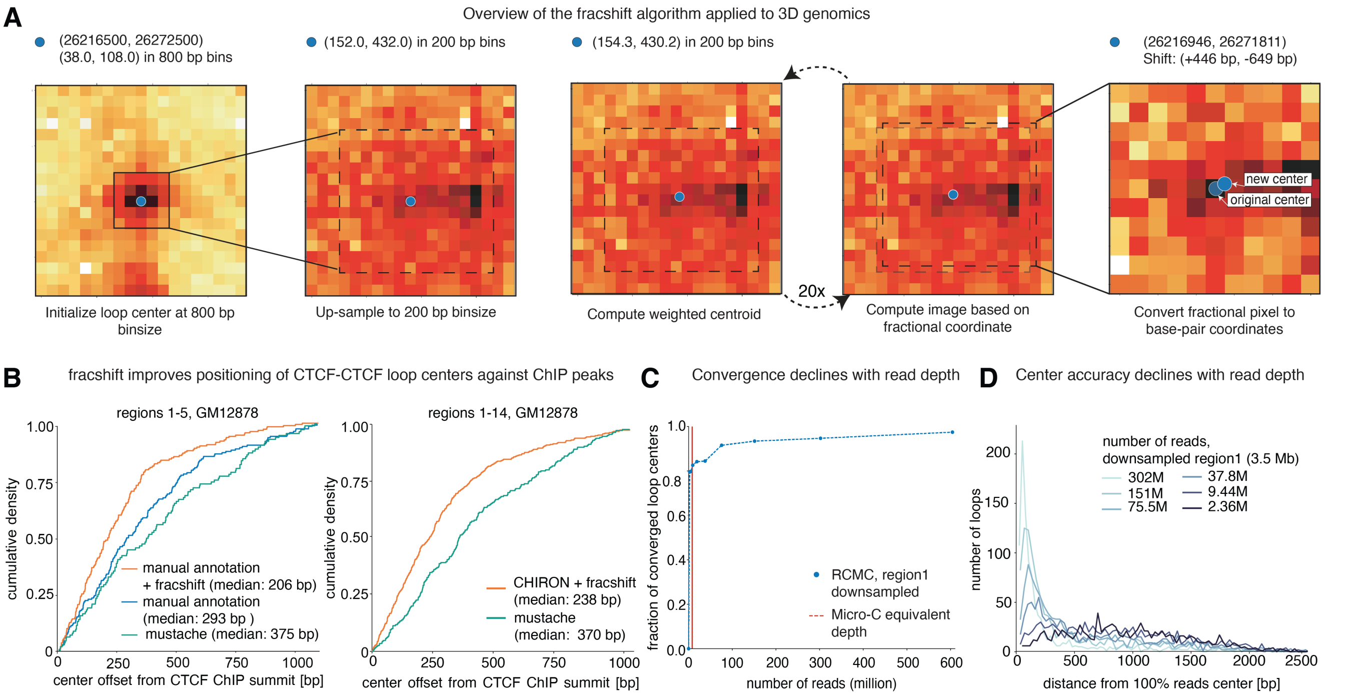


**Supplementary Figure 6: Sub-pixel localization of loop centers** (A) Implementation of the fracshift algorithm[15,16] for 3D genomic contact maps. Initial high-confidence centers are assigned at a larger binsize, then refined at a smaller target binsize. Fractional centers are computed by 20 iterations of intensity-weighted centroid estimation. (B) Validation of fracshift performance by computing the distance between CTCF-CTCF loop anchor positions and CTCF ChIP peak summits. (C) Number of loops for which the fracshift algorithm converges as a function of read depth (computed by downsampling GM12878 data from region1). (D) Distribution of loop center distances of fracshift-positioned centers in downsampled data from fracshift-positioned centers in the original data (GM12878, region1).

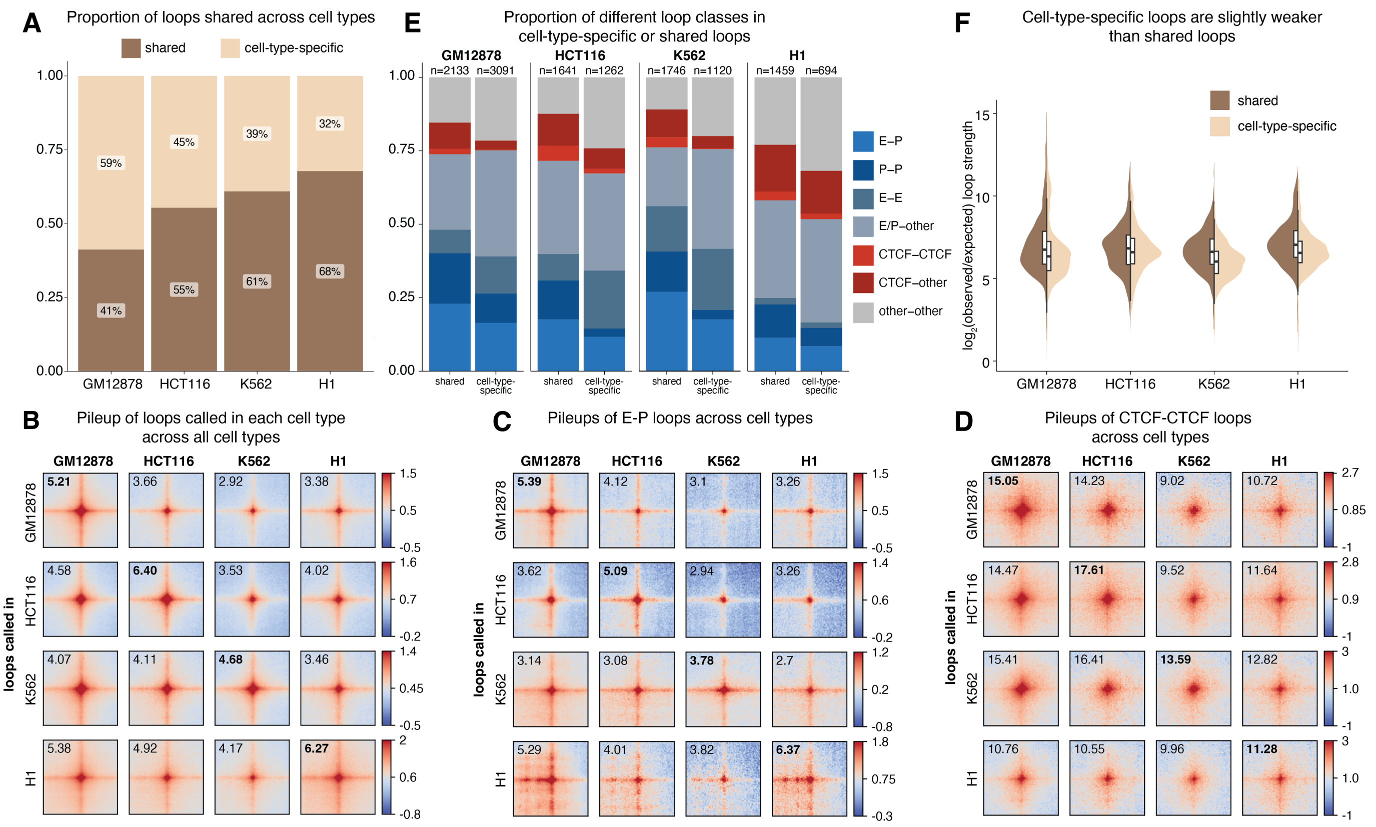


**Supplementary Figure 7: Characterization of looping interactions identified in RCMC.** (A) Proportion of loops that are shared with at least one other cell type (shared) or only present in the respective cell type (cell-type-specific). (B) Pileup of all loops called in each cell type (y-axis) on RCMC maps of all cell types. Values represent average loop strength. The enrichment of loops called in their respective cell types is highlighted with bold values. (C)-(D) Pileup of E-P loops (C) or CTCF loops (D) called in each cell type (y-axis) on RCMC maps of all cell types. Values represent average loop strength. The enrichment of loops called in their respective cell types is highlighted with bold values. (E) Distribution of loop classes for cell-type-specific or shared loops. (F) Distribution of loop strengths of cell-type-specific loops or loops that are shared with at least one other cell type.

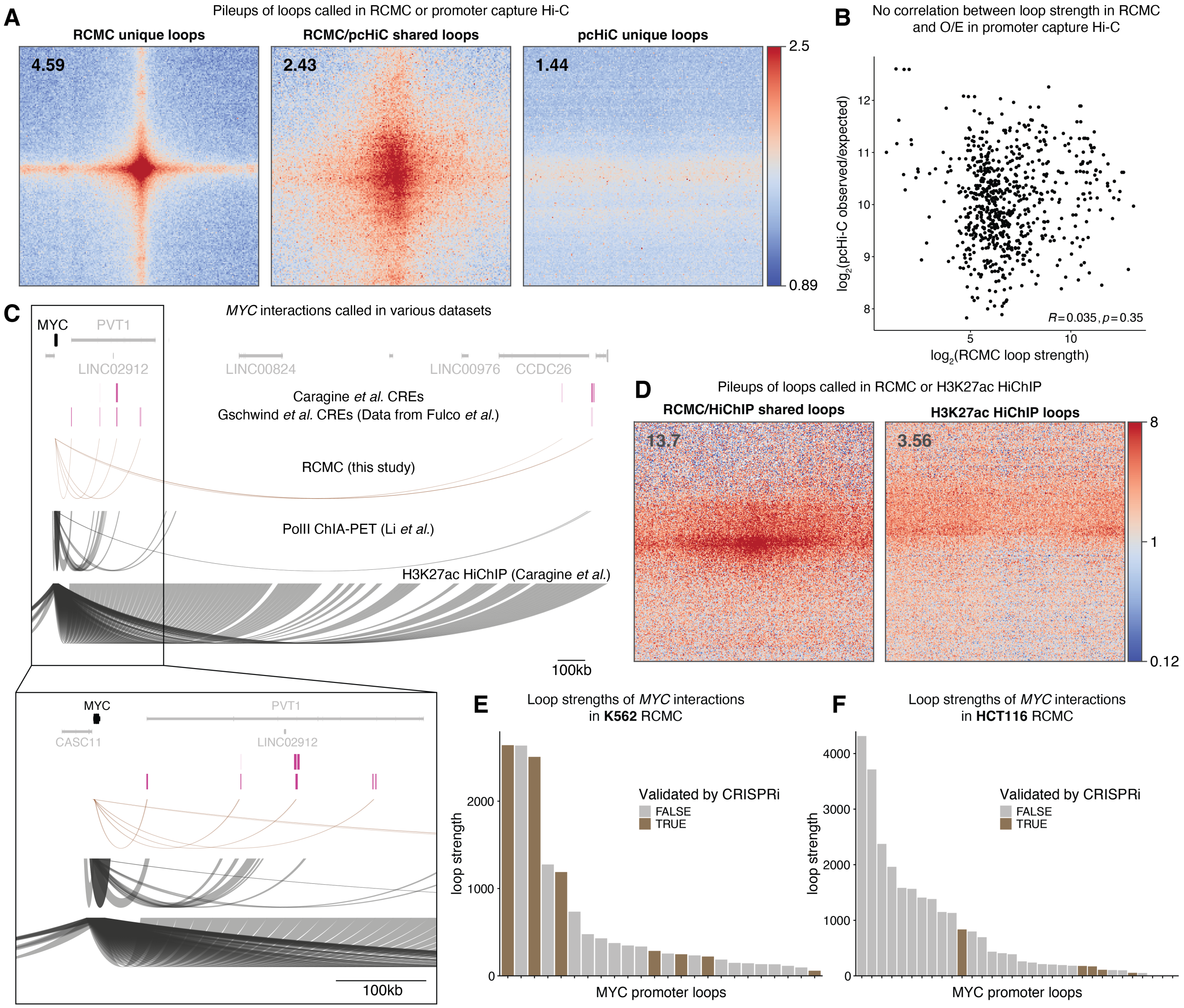


**Supplementary Figure 8: RCMC outperforms other capture-based methods at detecting functional loops.** (A) Pileup of loops unique to RCMC, promoter capture Hi-C (pcHi-C)[17] or shared between the two. Values represent average loop strength. (B) Loop strength comparison between RCMC and promoter-capture Hi-C for loops that are shared, where each point represents a shared loop. Pearson's correlation and significance are shown in the bottom right. (C) Comparison of loops interacting with *MYC* in RCMC, PolII ChIA-PET[24] or H3K27ac HiChIP[27] in K562. Arcs connect the two points that are looped, and the width of the arcs at the ends correspond to the bin sizes loops were called at. (D) Pileup of loops unique to H3K27ac HiChIP[27] or shared between RCMC and HiChIP. Values represent average loop strengths. (E)-(F) Loop strength of all loops interacting with *MYC* in K562 (E) or HCT116 (F) sorted by descending strength. Colored bars represent loops with enhancers that have been validated by at least one CRISPRi dataset.

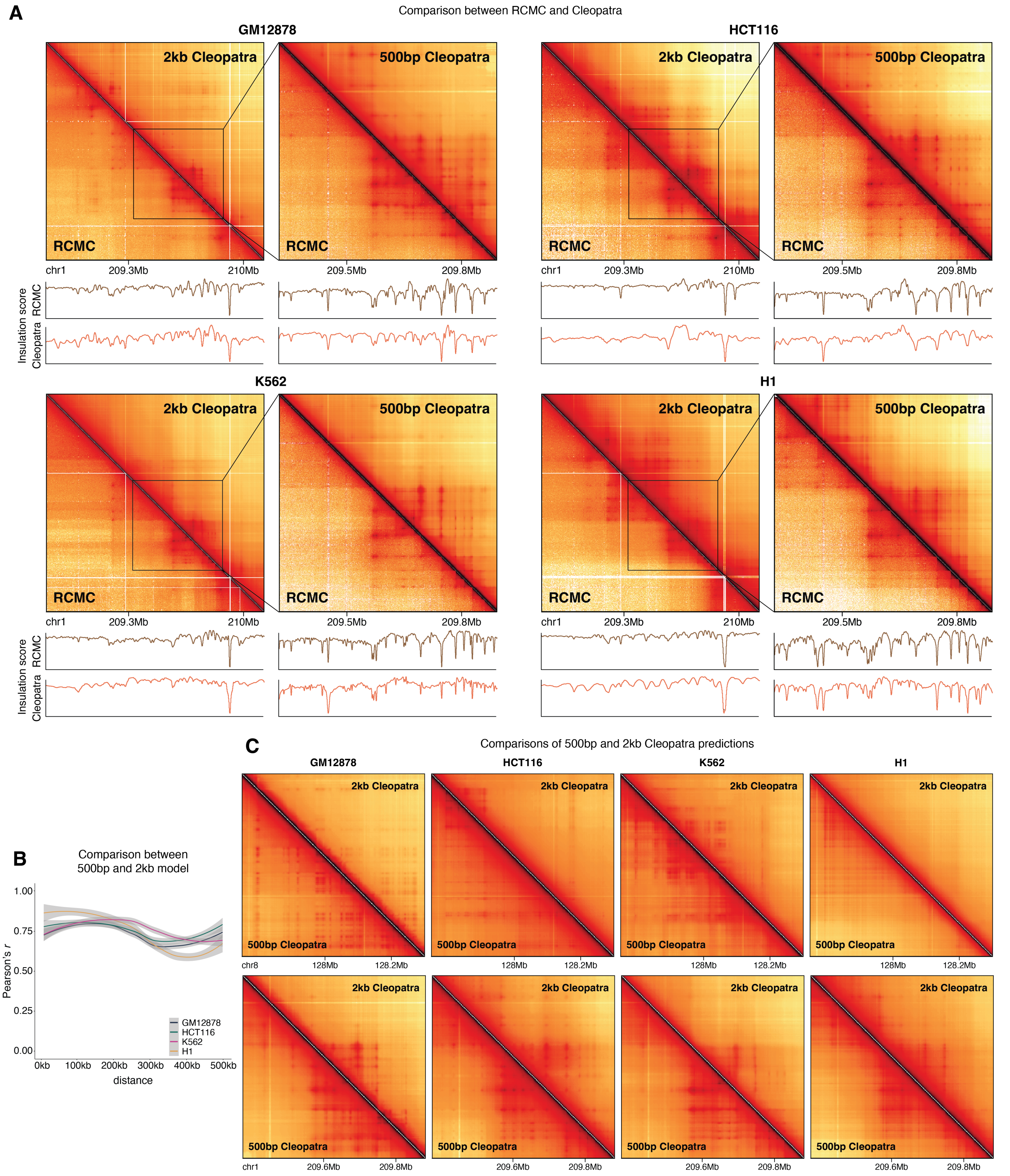


**Supplementary Figure 9: Cleopatra accurately predicts fine-scale 3D genome structures at small bin sizes.** (A) Cleopatra predictions in representative holdout region (region6) in four cell types at 2kb (left) and 500bp (right) bin sizes. Insulation scores for the corresponding regions from RCMC and Cleopatra are plotted below the maps. (B) Distance-stratified pearson's correlations between 500bp and 2kb Cleopatra models. (C) Example region showing that 500bp and 2kb Cleopatra predictions agree well with each other in holdout regions.

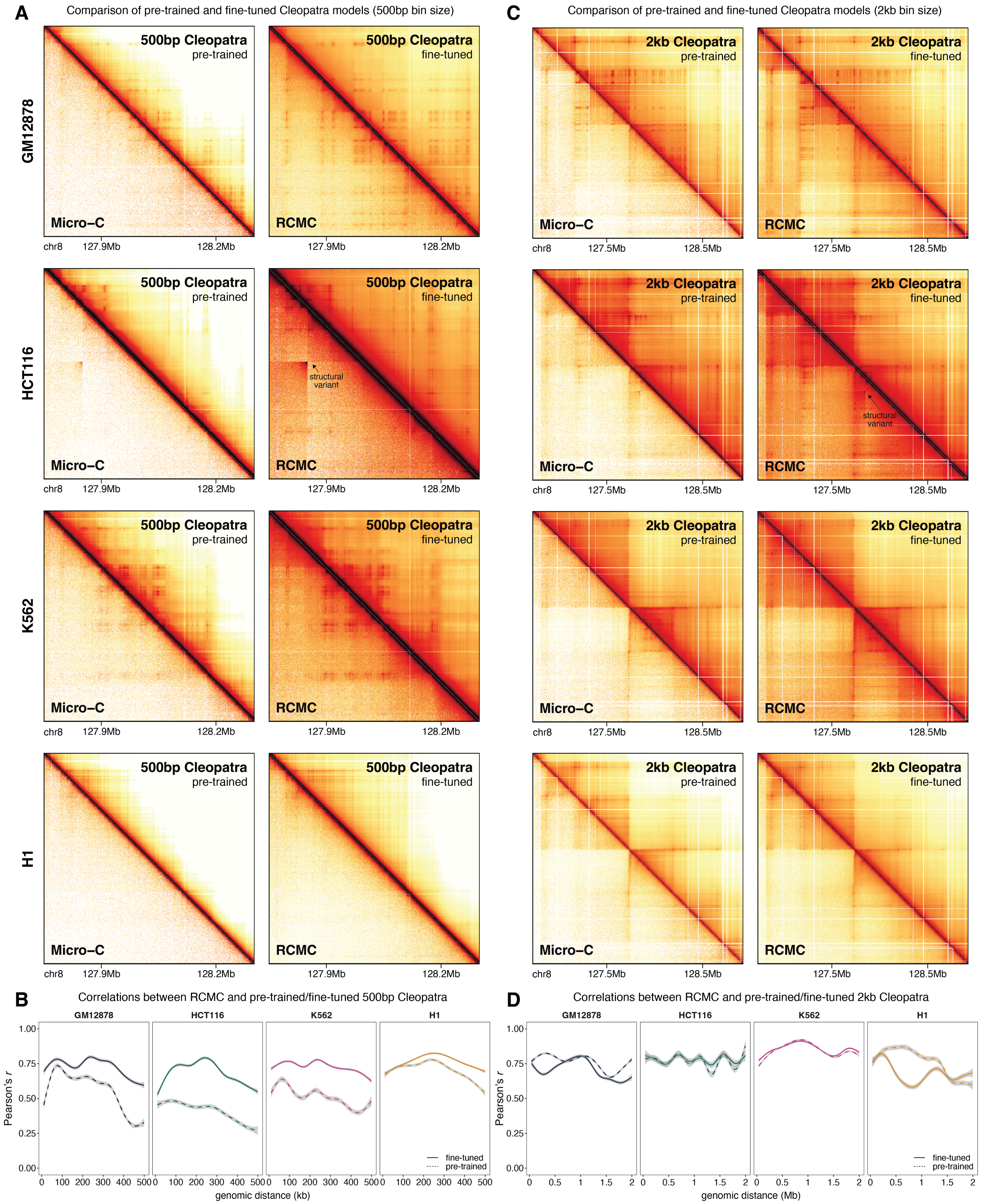


**Supplementary Figure 10: Cleopatra predicts RCMC-specific structures that are not visible in Micro-C at sub-kilobase bin sizes.** (A) Example of 500bp Cleopatra predictions from pre-training and fine-tuning at 500bp bin size. (B) Distance-stratified pearson's correlations between RCMC and pre-trained or fine-tuned Cleopatra in holdout regions. Shaded area represents 0.95 confidence interval from loess smoothing across 3 regions. (C) Example of 2kb Cleopatra predictions from pre-training and fine-tuning at 2kb bin size. (B) Distance-stratified pearson's correlations between RCMC and pre-trained or fine-tuned Cleopatra in holdout regions. Shaded area represents 0.95 confidence interval from loess smoothing across 3 regions.

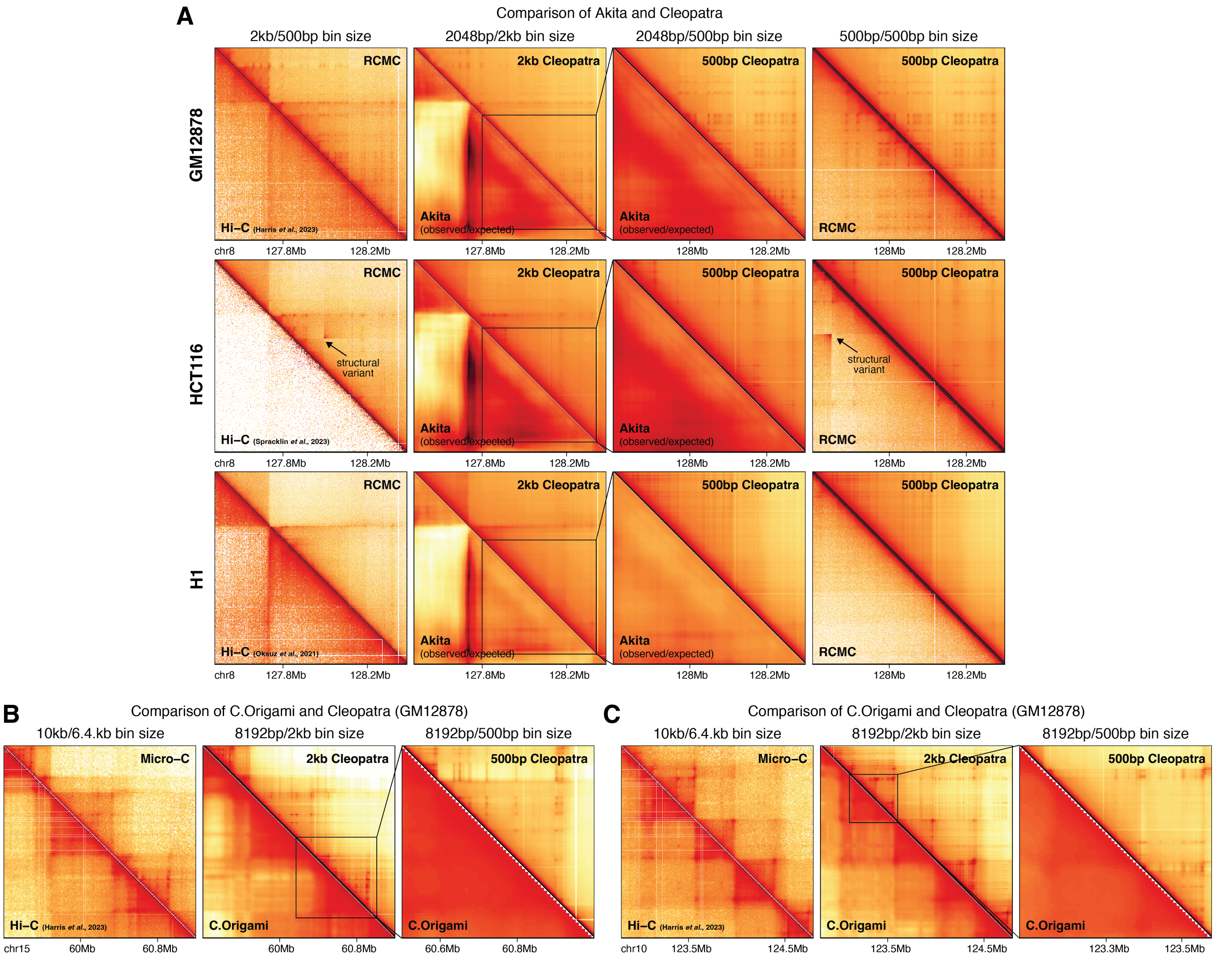


**Supplementary Figure 11: Cleopatra outperforms previous state-of-the-art 3D genome prediction models.** (A) Comparison of Akita V2[35] with Cleopatra at indicated bin sizes. This region is held out in both Akita and Cleopatra (region6). Note that Akita predictions are observed/expected, while Cleopatra predictions here are observed. While the comparison is not direct, it still highlights the fact that only Cleopatra can predict fine-scale structures. (B)-(C) Comparison of C.Origami[31] with Cleopatra at the indicated bin sizes. These regions were held out in C.Origami. No RCMC data is available, but Cleopatra predicts structures that are not present in Micro-C or Hi-C data. Hi-C data for GM12878 is from Harris *et al.*[53], HCT116 from Spracklin *et al.*[54] and H1 from Akgol Oksuz *et al.*[55].

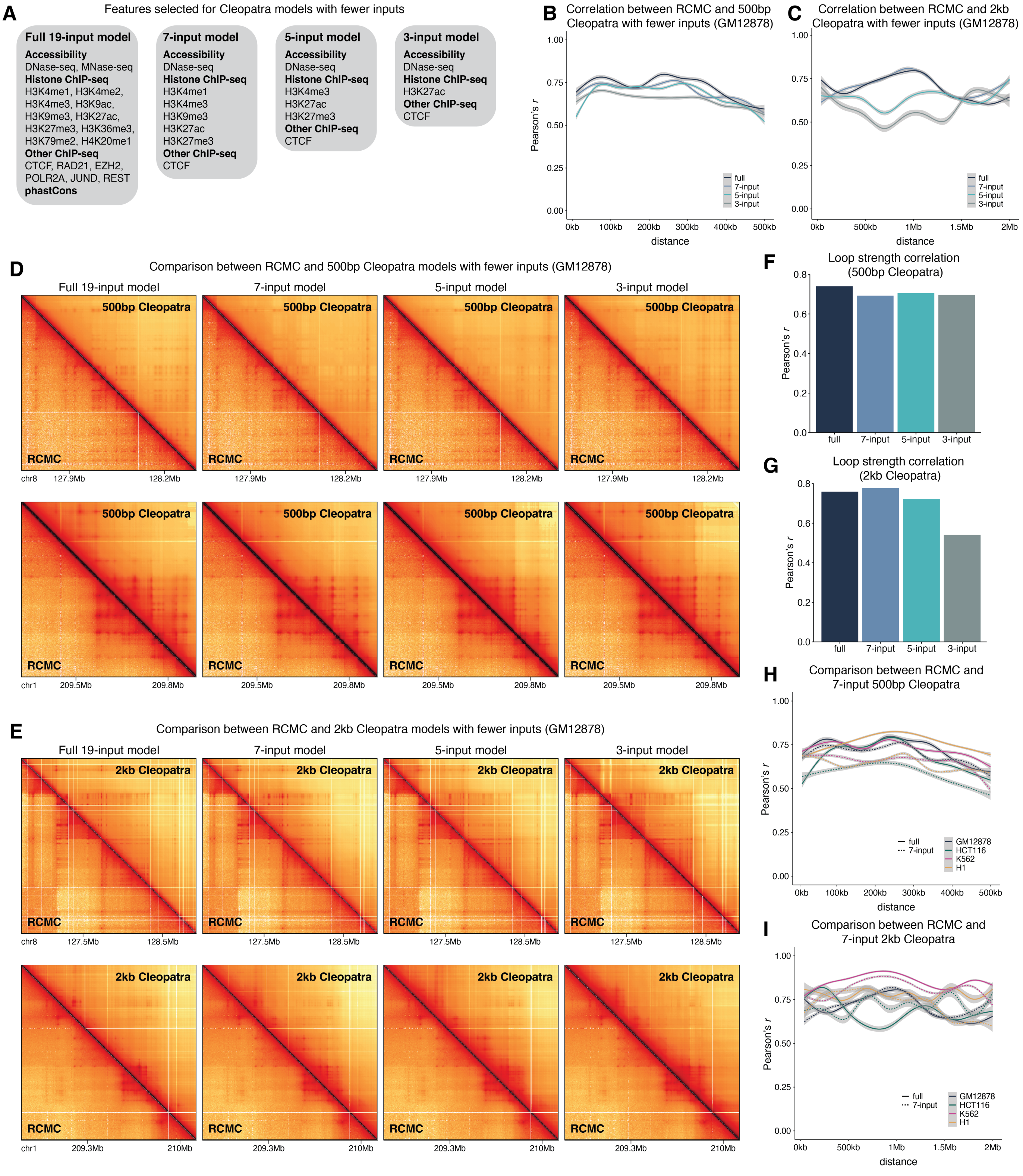


**Supplementary Figure 12: Cleopatra performance slightly decreases with fewer numbers of inputs.** (A) Features selected to train Cleopatra models with fewer inputs. (B)-(C) Distance-stratified pearson's correlations between RCMC and Cleopatra models with fewer inputs (500bp (B) and 2kb (C)) for GM12878. (D)-(E) Representative examples of GM12878 Cleopatra predictions with fewer inputs in two holdout regions (top: region6, bottom: region4). (F)-(G) Loop strength correlation between RCMC and Cleopatra models with fewer inputs in GM12878 (500bp (F) and 2kb (G)) (H)-(I) Distance-stratified pearson's correlations between RCMC and 7-input Cleopatra models for all cell types (500bp (H) and 2kb (I)). Shaded area represents 0.95 confidence interval from loess smoothing across 3 regions.

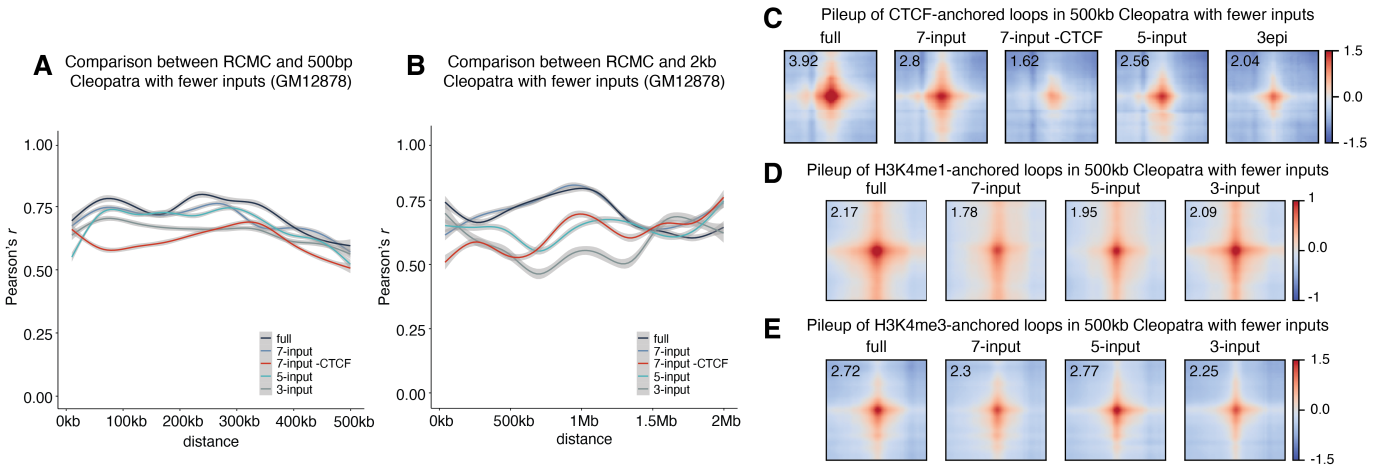


**Supplementary Figure 13: Contribution of features to GM12878 Cleopatra models.** (A)-(B) Distance-stratified pearson's correlations between RCMC and Cleopatra models with fewer inputs (500bp (A) and 2kb (B)). Shaded area represents 0.95 confidence interval from loess smoothing across three regions. (C) Pileup of CTCF-anchored loops (CTCF on both anchors) in Cleopatra models with fewer inputs. (D)-(E) Pileups of H3K4me1- (D) or H3K4me3- (E) anchored loops in 500kb Cleopatra models with fewer inputs. Numbers in all pileups represent average loop strengths.

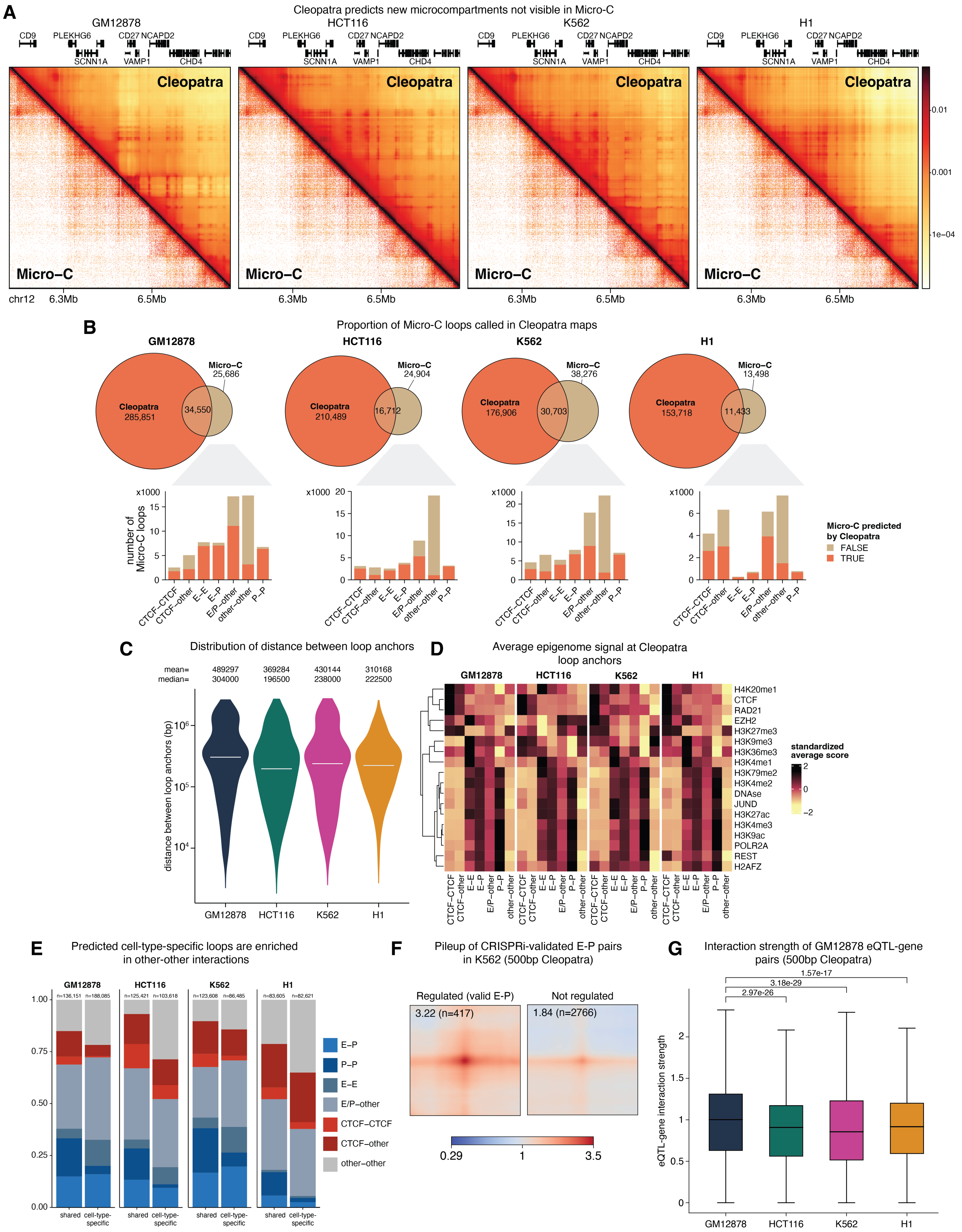
A
Cleopatra predicts new microcompartments not visible in Micro-C
GM12878
HCT116
K562
H1
CD9
PLEKHG6
CD27 NCAPD2
SCNN1A
VAMP1
CHD4
Cleopatra
Micro–C
chr12
6.3Mb
6.5Mb
0.01
0.001
1e–04
B
Proportion of Micro-C loops called in Cleopatra maps
GM12878
Micro–C 25,686
Cleopatra 285,851
34,550
HCT116
Micro–C 24,904
Cleopatra 210,489
16,712
K562
Micro–C 38,276
Cleopatra 176,906
30,703
H1
Micro–C 13,498
Cleopatra 153,718
11,433
x1000
number of Micro-C loops
CTCF–CTCF
CTCF–other
E–E
E–P
E/P–other
other–other
P–P
Micro-C predicted by Cleopatra
FALSE
TRUE
C
Distribution of distance between loop anchors
mean= 489297 369284 430144 310168
median= 304000 196500 238000 222500
distance between loop anchors (bp)
GM12878
HCT116
K562
H1
D
Average epigenome signal at Cleopatra loop anchors
H4K20me1
CTCF
RAD21
EZH2
H3K27me3
H3K9me3
H3K36me3
H3K4me1
H3K79me2
H3K4me2
DNAse
JUND
H3K27ac
H3K4me3
H3K9ac
POLR2A
REST
H2AFZ
standardized average score
E
Predicted cell-type-specific loops are enriched in other-other interactions
n=136,151 n=188,085
n=125,421 n=103,618
n=123,608 n=86,485
n=83,605 n=82,621
shared
cell-type-specific
E–P
P–P
E–E
E/P–other
CTCF–CTCF
CTCF–other
other–other
F
Pileup of CRISPRi-validated E-P pairs in K562 (500bp Cleopatra)
Regulated (valid E-P)
Not regulated
3.22 (n=417)
1.84 (n=2766)
0.29
1
3.5
G
Interaction strength of GM12878 eQTL-gene pairs (500bp Cleopatra)
1.57e-17
3.18e-29
2.97e-26
eQTL-gene interaction strength
GM12878
HCT116
K562
H1

**Supplementary Figure 14:** (Figure on previous page) **Cleopatra reveals new looping interactions.** (A) Representative new microcompartment predicted by Cleopatra (500bp model) that was previously not visible in Micro-C. (B) Proportion of Micro-C loops predicted by Cleopatra for each loop class. (C) Distribution of loop size in Cleopatra loops. White line indicates median value for each cell type. (D) Mean signals of various ChIP signals for each loop class. Coverage of each signal was calculated for each 1kb loop anchor, then averaged across all loop anchors in a given loop class. The signals were then standardized across the row to ensure that all signals are on the same scale. (E) Distribution of loop classes for cell-type-specific or shared loops. (F) Pileups of Regulated/Not regulated loops in 500bp Cleopatra (K562) taken from the harmonized CRISPRi data in Gschwind *et al.*[50]. (G) Interaction scores of GM12878 eQTL-gene pairs in all cell types. The box extends from the first quartile to the third quartile of the data, with a line at the median. The whiskers extend from the box to the farthest data point lying within 1.5x the inter-quartile range from the box. p-values were calculated using a two-sided Mann-Whitney U test.

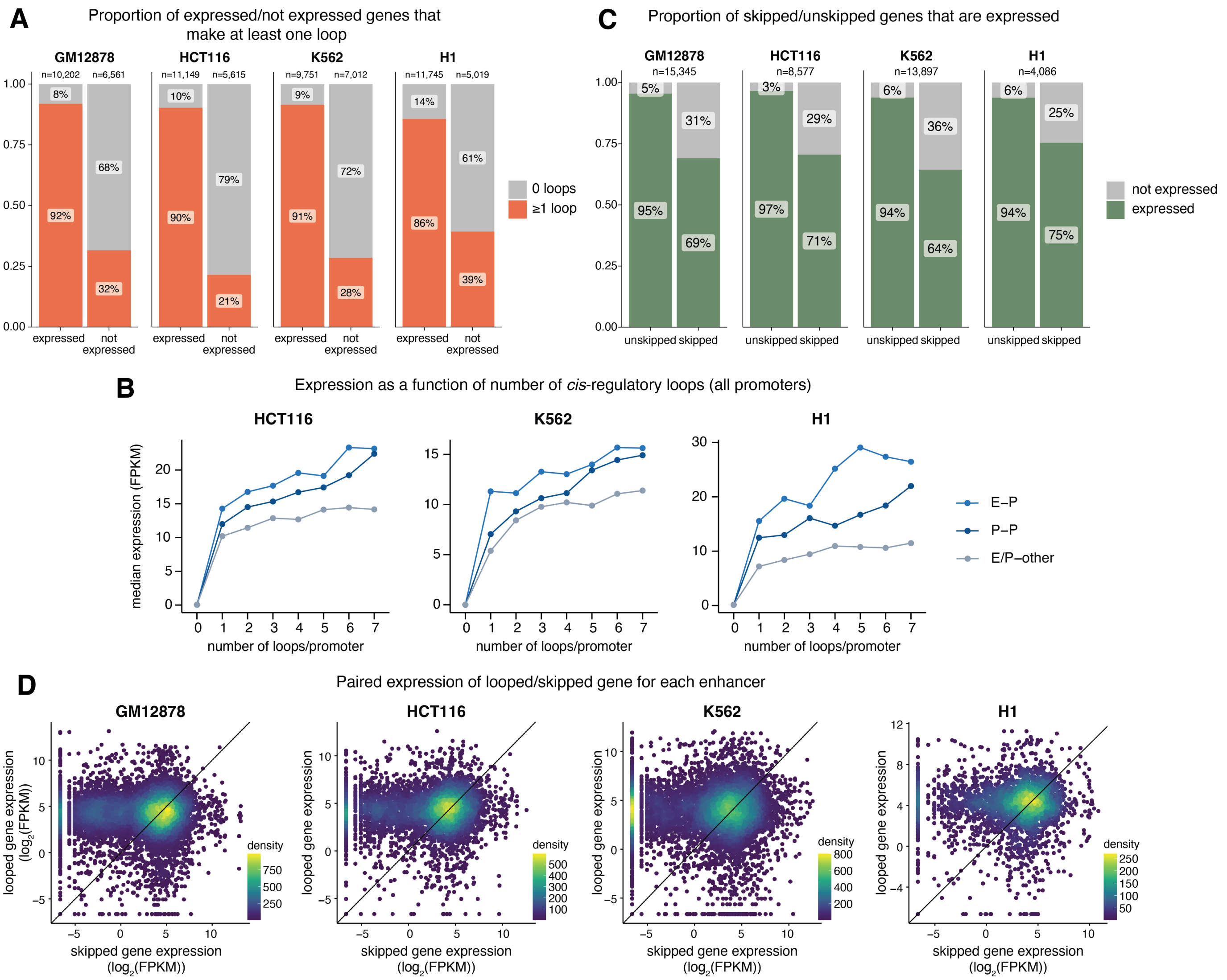


**Supplementary Figure 15: Relationship between looping interactions and gene expression revealed by Cleopatra.** (A) Proportion of genes make no loops or at least one loop. Gene promoters are defined as 1kb upstream and 500bp downstream of the transcription start site using GENCODE v29 annotations[56]. (B) Median gene expression of promoters with increasing numbers of E-P/P-P/P-other loops in HCT116, K562 and H1. Only promoters with $\leq$ 7 loops are shown here. Expression is calculated as the median FPKM of all promoters, including non-expressed promoters. (C) Proportion of unskipped/skipped genes that are not expressed. "Expressed" genes are defined as having FPKM >1 in RNA-seq data. (D) Scatterplot of paired skipped/looped gene expression per enhancer. For all enhancers that skip genes, the expression of the skipped gene is plotted against the further gene it is looped to. The color indicates the density of genes, and line indicates y=x.

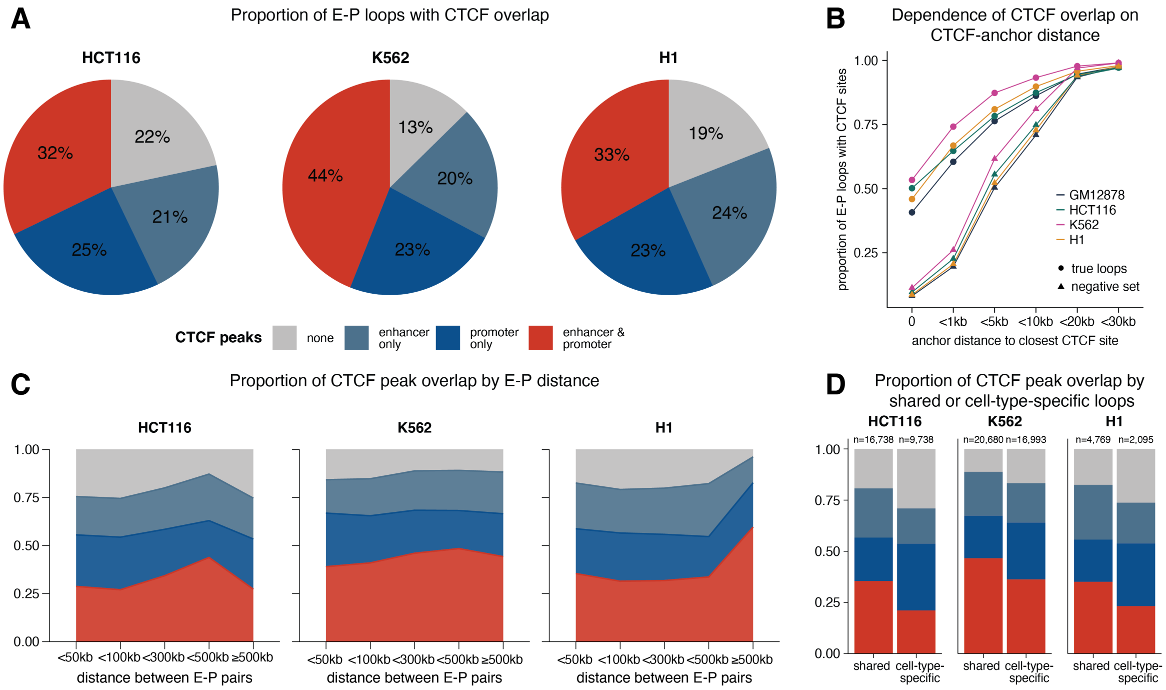


**Supplementary Figure 16: CTCF appears to facilitate looping interactions between enhancers and promoters.** (A) Proportion of E-P loops with CTCF binding at both the enhancer and promoter or enhancer/promoter only, using a 5kb window size around the midpoint. (B) Fraction of E-P loops that also contain CTCF sites depends on the distance threshold used to define CTCF overlap. The distance refers to distance betwen the midpoint of the enhancer or promoter and the apex of the CTCF peak. The true loops refers to Cleopatra-predicted E-P loops, while the negative set are the same loops shifted by 20kb downstream. (C) Proportion of E-P loops with CTCF binding on either anchor stratified by distance between the enhancer and promoter. (D) Proportion of E-P loops with CTCF binding on either anchor grouped by whether the E-P interaction is cell-type-specific or shared with at least one other cell type.

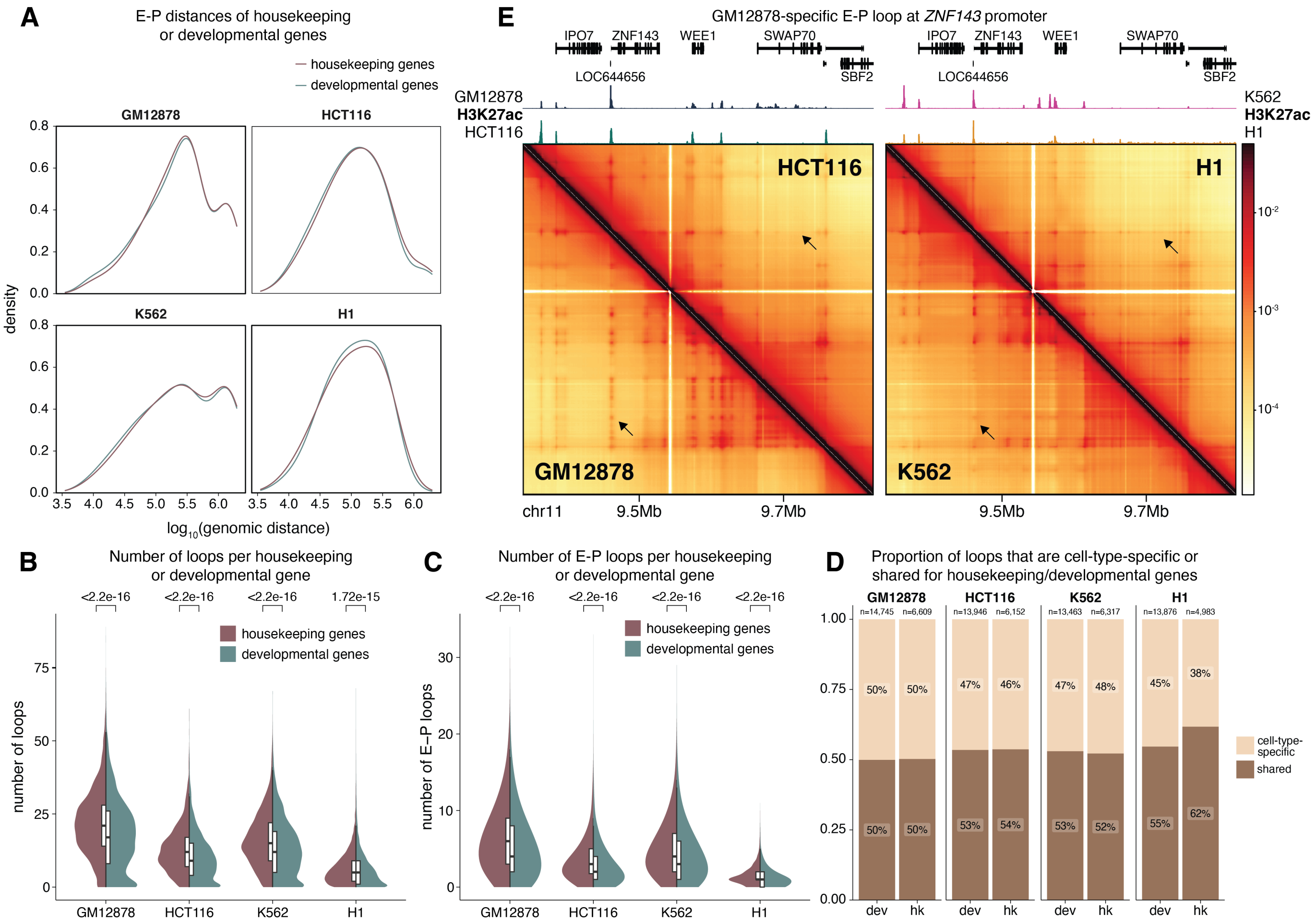


**Supplementary Figure 17: Similarities and differences between housekeeping and developmental genes revealed by Cleopatra.** Housekeeping and developmental gene annotations were obtained from Eisenberg & Levanon[57]. Only expressed genes (FPKM $\geq$1 were considered in the following analyses. (A) Distribution of E-P distance for housekeping or developmental genes. (B)-(C) Distribution of the number of all loops (B) or E-P loops (C) formed by each housekeeping or developmental promoter. p-values were calculated by a two-sided Mann-Whitney U test between housekeeping and developmental genes for each cell type. (D) Proportion of cell-type-specific or shared loops for housekeeping or developmental genes. (E) Example of cell-type-specific loop formed at the promoter of housekeeping gene *ZNF143*, indicated by the arrow. Arrows in other cell types point to corresponding region where no loop is observed.

#### Supplementary Tables

Table S1: Overview of all regions, number of replicates and number of reads obtained.
Table S2: Motifs identified in cell-type-specific fine-scale boundaries.
Table S3: Regions covered by region capture probes (all three sets combined).
Table S4: Other data sources used in the manuscript but not in Cleopatra training.
Table S5: All epigenomic data sources used for training Cleopatra.
Table S6: Loops identified across all RCMC data with loop class annotations and whether the loop is cell-type-specific.
Table S7: Loops called by Mustache identified in Cleopatra with loop class annotations and whether the loop is cell-type-specific. The column "called_from" indicates whether the loop was identified in 500kb or 2Mb Cleopatra.
